# Age-related Increase in Locus Coeruleus Activity and Connectivity with Prefrontal Cortex during Ambiguity Processing

**DOI:** 10.1101/2024.12.21.629084

**Authors:** Arjun Dave, Shuer Ye, Leona Rahel Bätz, Xiaqing Lan, Heidi I.L. Jacobs, Maryam Ziaei

## Abstract

Interpreting ambiguous environmental cues, like facial expressions, becomes increasingly challenging with age, especially as cognitive resources decline. Managing these challenges requires adaptive neural mechanisms that are essential for maintaining mental well-being. The locus coeruleus (LC), the brain’s main norepinephrine source, regulates attention, arousal, and stress response. With extensive cortical connections, the LC supports adapting to cognitive demands and resolving conflicting cues from environment, particularly in later life. Previous research suggests that LC interacts with the prefrontal cortex (PFC) during high-conflict tasks. However, whether LC activity and its connectivity with the PFC support emotional ambiguity processing and contributes to emotional well-being in healthy aging remains unclear. To address this gap, we used 7T-MRI to examine LC function in 75 younger (25.8 ± 4.02years, 35females) and 69 older adults (71.3 ± 4.1 years, 35females) during facial-emotion-recognition task morphed with varying ambiguity: anchor (unambiguous happy or fearful), intermediate-ambiguity (30%happy-70%fearful and 40%happy-60%fearful expressions), and absolute-ambiguity (50%happy-fearful). Behaviorally, participants had longer response times and lower confidence during the absolute-ambiguity condition, while older adults perceived ambiguous faces as happy more frequently than younger adults. Neuroimaging results revealed older adults exhibited greater LC activity and enhanced connectivity with dorsolateral PFC (dlPFC) during absolute-ambiguity compared to younger adults. This heightened connectivity in older adults was linked to better task-independent self-reported mental well-being questionnaires and greater emotional resilience scores derived from principal component analysis. Overall, these findings suggest that greater LC activity supports managing cognitively demanding tasks, while enhanced LC-dlPFC connectivity promotes emotional well-being, highlighting this neural pathway’s role in healthy aging.

**Significance Statement:** Understanding how the brain adapts to cognitive and emotional demands with age is key to promoting healthy aging. This study examined whether the locus coeruleus (LC), a brain region critical for regulating attention and arousal, undergoes adaptive changes with age, especially during emotional ambiguity task. Using ultra-high-field imaging, we explored younger and older adults recognize facial expressions with varying ambiguity levels. Our findings indicated that compared to young, older adults showed heightened LC activity and LC-dorsolateral prefrontal cortex (dlPFC) connectivity when processing absolute-ambiguous facial expressions, with enhanced connectivity linked to improved mental well-being. These results suggest higher LC activity supports cognitive demands of ambiguity processing with LC-dlPFC connectivity promoting emotional well-being and resilience, offering insights into mechanisms underlying healthy aging.

## Introduction

Interpreting facial expressions is fundamental to everyday social interactions, yet often impaired in neurological and psychiatric conditions (Henry et al., 2016). However, not all facial expressions convey a definitive emotional message. Ambiguity, such as a smile paired with furrowed brows, poses significant challenges in interpreting emotional intent (Zeki, 2004; Wang et al., 2017). Resolving such ambiguity necessitates sustained attention, arousal, and cognitive effort (FeldmanHall et al., 2016) key processes essential for interpreting ambiguous signals and supporting mental well-being (Keller et al., 2019; Wolpe et al., 2024). While older adults show differences in processing negative facial expressions (Ruffman et al., 2008), encoding ambiguous facial expression demands even greater cognitive effort. To navigate this, older adults tend to rely on adaptive neural mechanisms to mitigate cognitive decline (Dexter & Ossmy, 2023), though these mechanisms remain largely unknown.

The locus coeruleus (LC), a small brainstem structure, regulates key functions such as attention, arousal, emotion, memory, and stress response (Poe et al., 2020). It operates via two distinct firing modes: phasic bursts—brief and stimulus-locked that enhances focused attention and task-specific processing; whereas tonic firing—sustained at low frequency supports arousal, behavioral flexibility, and exploration during rest or low-demand states (Bouret & Sara, 2005; Aston-Jones & Cohen, 2005).

Additionally, the LC shapes cognition through extensive projections to the prefrontal cortex (PFC; Chandler et al., 2014), particularly the dorsolateral PFC (dlPFC; Levinson et al., 2023), underpinning top-down regulatory mechanisms critical for resolving conflicting information (Grueschow et al., 2020) and maintaining emotional balance and well-being (Morris et al., 2020; Watanabe et al., 2024). In late adulthood, structural and functional changes in the LC and its connectivity with the PFC contribute to reduced attention (Lee et al., 2020), impaired episodic memory (Dahl et al., 2023), and distractibility (Song et al., 2021). However, preserved LC structural integrity is associated with superior episodic, working memory, and cognitive reserve (Clewett et al., 2016; Dahl et al., 2019). Notably, elevated LC activity in response to emotionally salient stimuli, has also been linked to greater resilience against cognitive decline in later life (Clewett et al., 2018; Prokopiou et al., 2022; Ludwig et al., 2024). Furthermore, the LC-prefrontal functional coupling supports inhibitory control among older adults (Tomassini et al., 2022), suggesting its role in compensatory adaptations during healthy aging. However, despite its significance in cognitive performance, the LC-prefrontal pathway’s specific role in resolving emotional ambiguity and promoting mental well-being in late adulthood remains unexplored. Advances in neuroimaging, particularly 7-Tesla Magnetic Resonance Imaging (7T-MRI) provide a powerful tool for delineating and characterizing the LC, offering insights into adaptive processes that may help mitigate age-related cognitive and emotional challenges.

This study addresses this gap by investigating age-related differences in LC activity and LC-PFC functional connectivity during emotional ambiguity processing, and their role in promoting emotional resilience and mental well-being in healthy aging. Using ultra-high field 7T-MRI, we examined responses to an emotion recognition task among healthy younger and older adults. The task required recognizing facial expressions ranging from unambiguous (100%-fearful or 100%-happy) to varying levels of ambiguity in morphed expressions (30% happy–70% fearful and 40% happy–60% fearful expressions, in either direction), including fully ambiguous expression (50%-fearful/50%-happy). Given age-related differences in LC activity during emotionally salient, novel, and cognitively demanding tasks (Prokopiou et al., 2022; Ludwig et al., 2024) and evidence that norepinephrine (NE) release during cognitively effortful tasks supports neural resilience and preserves cognitive function in late life (Wilson et al., 2013; Yu et al., 2015; Mather & Harley, 2016; Ciampa et al., 2022), we hypothesized that older adults would exhibit heightened LC responses to ambiguous expressions compared to younger adults. While evidence remains limited, based on the links between LC-prefrontal coupling with better inhibitory control (Tomassini et al., 2022), we anticipated stronger LC-PFC connectivity during emotional ambiguity processing among older adults, and expected this connectivity to relate with mental well-being, emphasizing this pathway’s relevance in promoting mental health in late adulthood.

## Materials and Methods

### Participants

To determine the minimum sample size required to detect a moderate-to-large age effect (f = 0.33) with 85% power (α = 0.01), we conducted a power analysis using G*Power version 3.1. The analysis indicated a requirement of a minimum of 70 participants per age group to achieve sufficient statistical power. Eighty younger and 75 older adults were recruited and scanned as part of the Trondheim Aging Brain Study (TABS) completed two sessions: one imaging and one behavioral-neuropsychological testing sessions. However, due to incomplete behavioral data sets, technical issues during data collection, and excessive head motion (mean framewise displacement ≥ 0.5 mm) in five young and six elderly participants, the final sample included a total of 144 participants, including 75 younger (mean age = 25.8 years, SD = 4.02; 35 females) and 69 older adults (mean age = 71.27 years, SD = 4.07; 35 females). Participants were recruited from the local community through paper flyers and online advertisements. Exclusion criteria included the presence of metallic implants, claustrophobia, or a history of neurological (e.g., neurodegenerative diseases), psychiatric disorders (major depression) or autism spectrum disorder. All participants were right-handed, had normal or corrected-to-normal vision, and reported normal hearing abilities. All older participants underwent cognitive screening using the Mini-Mental State Examination (MMSE; Folstein et al., 1975), conducted either in person or via phone using the 22-point Braztel-MMSE (Camozzato et al., 2010), and participants scored above the respective thresholds of 24 and 15, confirming lack of cognitive deficits. Specifically, MMSE scores ranged from 25 to 30 (M = 29.00, SD = 1.21), and telephonic Braztel-MMSE scores ranged from 17 to 22 (M = 21.34, SD = 0.99). Participants completed a range of self-report questionnaires and cognitive measures during the behavioral session (<u>Table S1</u>). All participants provided written informed consent and received compensation of 500 Norwegian Kroner in gift cards for their participation. This study was approved by the regional Ethics Committee.

### Experimental design

The experimental paradigm was developed and executed using MATLAB R2017b (The MathWorks) in conjunction with the Psychtoolbox extension (http://psychtoolbox.org). The task design was adapted from the Wang and colleagues work (2017). Facial expression stimuli were generated by morphing prototypical fearful and happy faces along a continuous gradient using the Delaunay tessellation function. This method involved selecting control points on the face manually, which were then used to guide the transformation. The pixel values and their spatial locations were interpolated through piecewise cubic-spline transformation, ensuring smooth transitions between facial features across the morphing continuum (Wang et al., 2017). The process allowed for the creation of highly realistic expressions that captured varying degrees of emotional intensity and ambiguity facilitating nuanced experimental manipulation of facial stimuli. The stimuli were created at seven distinct levels of emotional intensity, ranging from 100% fear to 100% happiness. Ambiguity levels spanned from 30% fearful - 70% happiness to 70% fearful - 30% happiness, with increments of 10%, including a 50% fearful - 50% happiness condition (Figure 1b). Low-level image properties, including Fourier amplitude spectra, mean luminance, contrast, and histogram specifications, were equalized using the SHINE toolbox (Willenbockel et al., 2010) to ensure perceptual visual quality. In total, a sample of four young and four elderly individuals (4 males and 4 females in each group) were sourced from the FACES database (Ebner et al., 2010). Faces were matched for attractiveness and distinctiveness with no difference between males and females within each emotional categories [attractiveness: M _male happy_ = 46.29 (SD = 7.90), M _female happy_ = 47.60 (SD = 7.15), M _male fearful_ = 28.83 (SD = 7.84), M _female fearful_ = 30.73 (SD = 8.62; Distinctiveness: M _male happy_ = 45.45 (SD = 3.72), M _female happy_ = 44.86 (SD = 4.20), M _male fearful_ = 42.35 (SD = 3.29), M _female fearful_ = 42.41 (SD = 5.09)]. An online pilot study was conducted to gauge responses to each facial expression and ambiguity levels created for this study. Seventy-four participants (40 females; M = 28.30 years, SD = 6.50 years) were shown morphed faces and asked to indicate emotional expressions as well as intensity of the facial expression on a 1 to 10 Likert scale. Findings of the pilot study indicated higher intensity rating for anchor images, such as happy and fearful (M = 7.57, SD = 0.80), followed by intermediate ambiguity (M = 5.86, SD = 1.05) and ambiguous faces (M = 5.02, SD = 0.83), confirming the effect of interest for ambiguity processing from the FACES dataset.

**Figure 1:**
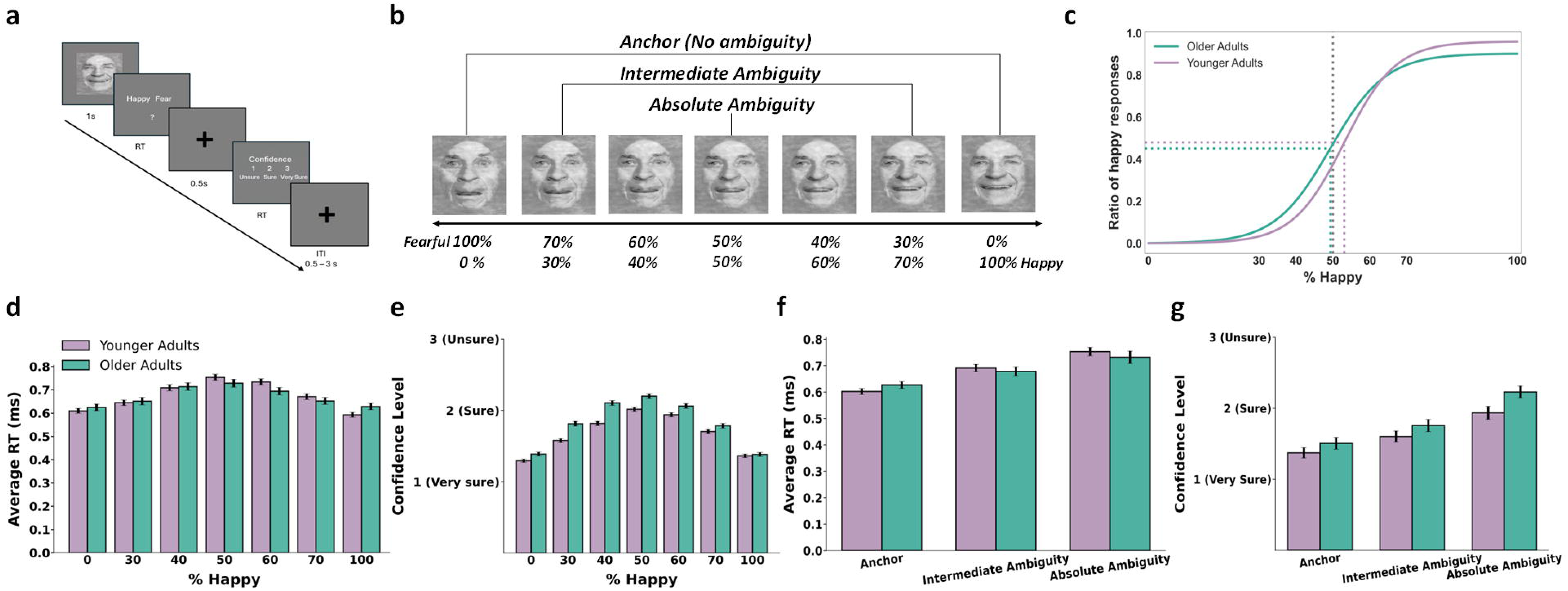
Experimental task and behavioral results. panel (a) shows an example of experimental Trial. A face was displayed for 1 second, after which participants were prompted to identify the facial emotion as either fearful or happy. Following a 500-millisecond cross, they were asked to rate their confidence by selecting ‘1’ for very sure, ‘2’ for sure, or ‘3’ for unsure. Panel (b) shows sample stimuli of one male ranging from 0% fear to 100% happy. (c-g) Behavioral results. Panel (c) depicts the group average of psychometric curves of both young and older adults showing the proportion of trials judged as happy normalized to 0 and 1 as a function of morph levels (ranging from 0% fearful to 100% happy). Panel (d) and (f) show response time (RT) comparison as a function of % happy and combined ambiguity level between younger and older subjects, respectively. Panels (e) and (g) depict confidence level comparison as a function of % happy and combined ambiguity level between younger and older subjects, respectively. Error bar denotes one standard error of mean (s.e.m) across subjects/sessions. Images of faces were derived from the FACES database (Max Planck Institute for Human Development). Used with permission under the terms of the database’s copyright policy.

The task design (Figure 1a) was event-related where a face was presented for 1 second, followed by a question asking participants to identify the emotional expression of the face. Participants indicated their decisions (fearful or happy) by pressing a button within 2 seconds followed by a confidence rating block. Participants rated their confidence level by pressing buttons: 1 for “very sure,” 2 for “sure,” or 3 for “unsure” within 2 seconds. Following confidence response rating, a cross was presented as the inter-trial intervals, which was jittered randomly, with a uniform distribution between 1 to 3.5 seconds within each run. Each participant completed a total of 168 trials with a total time of ∼21 minutes across three runs. The order of runs was counterbalanced between participants and order of faces within each run was pseudorandomized, ensuring no more than two consecutive faces shared the same emotional expression, identity, or age category. Behavioral responses were collected through an MR-compatible button box.

For subsequent analyses, the conditions were grouped based on their level of ambiguity: 100% fearful and 100% happy facial expressions were categorized as an anchor condition, the 50% happy - 50% fear morphed face as an absolute-ambiguous condition, and the remaining morphed facial expression (30, 40, 60, and 70) as an intermediate-ambiguity condition. This grouping was chosen to assess whether age-related differences in behavioral and neural responses are influenced by varying levels of ambiguity.

### Cognitive and emotional well-being measurements

As part of the TABS, participants completed a comprehensive series of behavioral and neuropsychological tests outside the scanner, designed to evaluate executive functioning, empathy, theory of mind, and overall mental well-being. The behavioral testing session was completed within two to four days following the imaging session followed by a 7-days daily sampling method, which is not reported here. Descriptives and inferential statistics of cognitive and emotional measures are reported in <u>Table S1</u>. Executive control functioning was assessed using a battery of tests, including the Stroop test (Jensen & Rohwer, 1966), the Trail Making Test (both A and B parts, Reitan & Wolfson, 2004), and the Phonemic and Semantic Verbal Fluency Task (Newcombe, 1969). Theory of Mind and empathy were measured using the Reading the Mind in the Eyes Task (RMET; Baron-Cohen et al., 2001) and the Interpersonal Reactivity Index (IRI; Davis, 1980), respectively. The abbreviated version of Raven’s Progressive Matrices (Raven, 1940) was used for measuring general intelligence. Emotional and mental well-being were evaluated using several validated questionnaires, including the Depression, Anxiety, and Stress Scale (DASS-21; Lovibond & Lovibond, 1995), the Difficulties in Emotion Regulation Scale (DERS; Gratz & Roemer, 2004), the General Health Questionnaire (GHQ-28; Goldberg, 1979), the Hospital Anxiety and Depression Scale (HADS; Zigmond & Snaith, 1983), the Connor-Davidson Resilience Scale (CD-RISC; Connor & Davidson, 2003), the Intolerance of Uncertainty Scale (IUS; Freeston et al., 1994; Buhr & Dugas., 2002), the Perceived Stress Scale (PSS; Cohen et al., 1983) and the State-Trait Anxiety Inventory (STAI; Spielberger, 1983). As shown in <u>Table</u> <u>S1</u>, younger and older adults had comparable scores in cognitive and emotional empathy scores, however, older adults reported better mental well-being and higher emotional resilience, with lower scores on self-reported measures of depression, stress, and anxiety, and performed better on the social cognitive tasks such as the Reading the Mind in the Eyes Test (RMET), compared to younger adults. In contrast, younger adults outperformed older adults in working memory, IQ, verbal fluency, and trail-making tests (all *p*s < 0.05).

### Emotional well-being factor analysis

Participants were assessed on multiple emotional and cognitive measures using established scales, and Principal Component Analysis (PCA) was applied to reduce data dimensionality for both cognitive and emotional assessments separately. This resulted in the identification of two primary indices: the ‘Emotional Resilience Index’, which captures the multidimensional construct of emotional resilience and mental well-being, and the ‘Cognitive Function Index’, representing performance on cognitive tasks. Sampling adequacy for the Emotional Resilience Index was confirmed with a high Kaiser-Meyer-Olkin (KMO) value of 0.918, while Bartlett’s test of sphericity indicated strong correlations among variables (χ²_(142)_ = 105, *p* < 0.001) (Masullo et al., 2021). For the Cognitive Function Index, a moderate KMO value of 0.67 was observed, with Bartlett’s test similarly confirming strong correlations among variables (χ²_(142)_= 21, *p* < 0.001). PCA components were considered significant if their eigenvalues exceeded 1. The Emotional Resilience Index loaded on two significant components, explaining 63.94% of the variance in total, with the first component explaining 57.05% of variance. In contrast, the Cognitive Function Index loaded on three significant components, accounting for 72.69% of the total variance, with the first and second component explaining 37.26% and 20.06%, respectively. Composite scores for both indices were calculated as weighted sums of the factor scores, with weights determined by the variance explained by each factor (Antony & Rao, 2007). The Emotional Resilience Index reflects an individual’s ability to feel, manage and regulate emotional responses, along with adapting to uncertain situations. The Cognitive Function Index serves as a comprehensive measure of cognitive performance, encompassing key domains such as verbal ability (assessed through COWAT Phonemic and Semantic tasks), working memory (evaluated via forward and backward digit span tests), and inhibitory control (measured using the Stroop task). To enhance interpretability, scores for the Emotional Resilience Index were inverted, indicating higher values for better emotional resilience.

### Behavioral data analysis

Behavioral results are presented in Figure 1 for seven conditions as well as three grouped conditions. Response bias, response times (RT) and confidence rating were analyzed separately. Given that the ambiguity responses do not have any correct responses, we included all responses for behavioral analyses. For statistical analysis, a 2 (age group: young, old) by 3 (ambiguity level: anchor, intermediate, absolute-ambiguous) repeated-measures analysis of variance (ANOVA) was conducted, with RT and confidence rating as dependent variables. This analysis allowed for the evaluation of both between-group differences in age and ambiguity level, as well as the interaction between factors.

The responses to faces were analyzed using a logistic function to generate the smooth psychometric curves shown in Figure 1c similar to Wang et al. (2017):

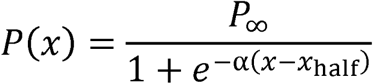

where *P* represents the percentage of trials where faces were judged as fearful, _X_ is the morph level, *P*_∞_ is the maximum value as _X_ approaches infinity, _Xhalf_ is the curve’s midpoint, and α indicates curve steepness. These parameters (*P*_∞_, *X*_half_ and α) were fitted to each subject’s data from the emotion perception task. Flatter curves (lower α) indicate reduced sensitivity to emotion changes, while steeper curves (higher α) indicate greater sensitivity.

### MRI acquisition and analyses

#### Image acquisition

MR scans were acquired on a 7T Siemens MAGNETOM Terra scanner with a 32-channel head coil at the Norwegian 7T MR Center at St. Olav’s Hospital. First, high-resolution anatomical T1-weighted (T1w) images were acquired using a MP2RAGE sequence (Marques et al., 2009; voxel size = 0.75 × 0.75 × 0.75 mm, with 224 slices, repetition time (TR) = 4300 ms, echo time (TE) = 1.99 ms, flip angles = 5°/6°, field-of-view (FOV) = 240 × 240 × 175 and slice thickness = 0.75 mm). To achieve high-resolution imaging of the LC, we employed a magnetization transfer-weighted turbo flash (MT-TFL) sequence (Priovoulos et al., 2018), which is sensitive to LC contrast and utilizes a multi-shot 3D readout (voxel size = 0.4 × 0.4 × 0.5 mm, with 60 slices, TR = 400 ms, TE = 2.55 ms, flip angle = 8°). The FOV for the MT-TFL sequence was positioned approximately perpendicular to the pons, encompassing the region between the inferior colliculus and the inferior border of the pons. Whole-brain functional images were collected using T2*-weighted echo-planar image sequence (90 interleaved slices, TR = 2s, TE = 35 ms, voxel size = 1.25 × 1.25 × 1.25 mm, slice thickness = 1.25mm, multiband factor = 2, flip angle = 80°, FOV = 200 × 200 × 112.5 mm).

#### Template creation and LC delineation

The procedure of the LC delineation is shown in Figure 2. The MT-TFL and whole-brain T1w images were utilized to create a study-specific template using the *antsMultivariateTemplateConstruction2.sh* function from Advanced Normalization Tools (ANTs). This process employed default parameters, with the exception that rigid-body registration was enabled. Prior to template construction of T1w, intensity non-uniformity was corrected using the *N4BiasFieldCorrection* method within ANTs (Tustison et al., 2010). To delineate LC structure, we followed the methodology outlined by Van Egroo et al. (2021, 2023; Figure 2). Briefly, individual MT-TFL images were intensity-normalized by dividing each image by the subject-specific mean intensity measured within a 10×10 voxel region of interest (ROI) located in the pontine tegmentum (PT). For each individual participant, the PT-ROI was consistently placed in the axial slice containing the highest intensity LC voxel identified by the first author (Figure 2b). By normalizing to the PT—an anatomical region adjacent to the LC with homogeneous tissue properties (Liu et al., 2017), we accounted for inter-subject variability in signal intensity, while preserving the relative contrast necessary for accurate LC localization (Tona et al., 2019). Subsequently, a study-specific template was constructed from these intensity-normalized MT-TFL images (Figure 2c) using the *antsMultivariateTemplateConstruction2* function in ANTs (Avants et al. 2010), with the transformation model and similarity metric set to greedy SyN and cross-correlation, respectively. The LC was manually delineated on the resulting template (Figure 2d, step 1) based on voxel intensities and anatomical characteristics using ITK-SNAP (by first author). This segmentation was repeated four weeks later by the first author, with only voxels identified in both sessions included in the final segmentation (Jacobs et al., 2020). To assess reliability, we calculated the Sørensen–Dice coefficient between the two independent segmentations, yielding values of 0.959 for the left LC mask and 0.962 for the right LC mask, indicating high consistency across sessions in the LC segmentation (Check LC Volume in Text S1).

**Figure 2:**
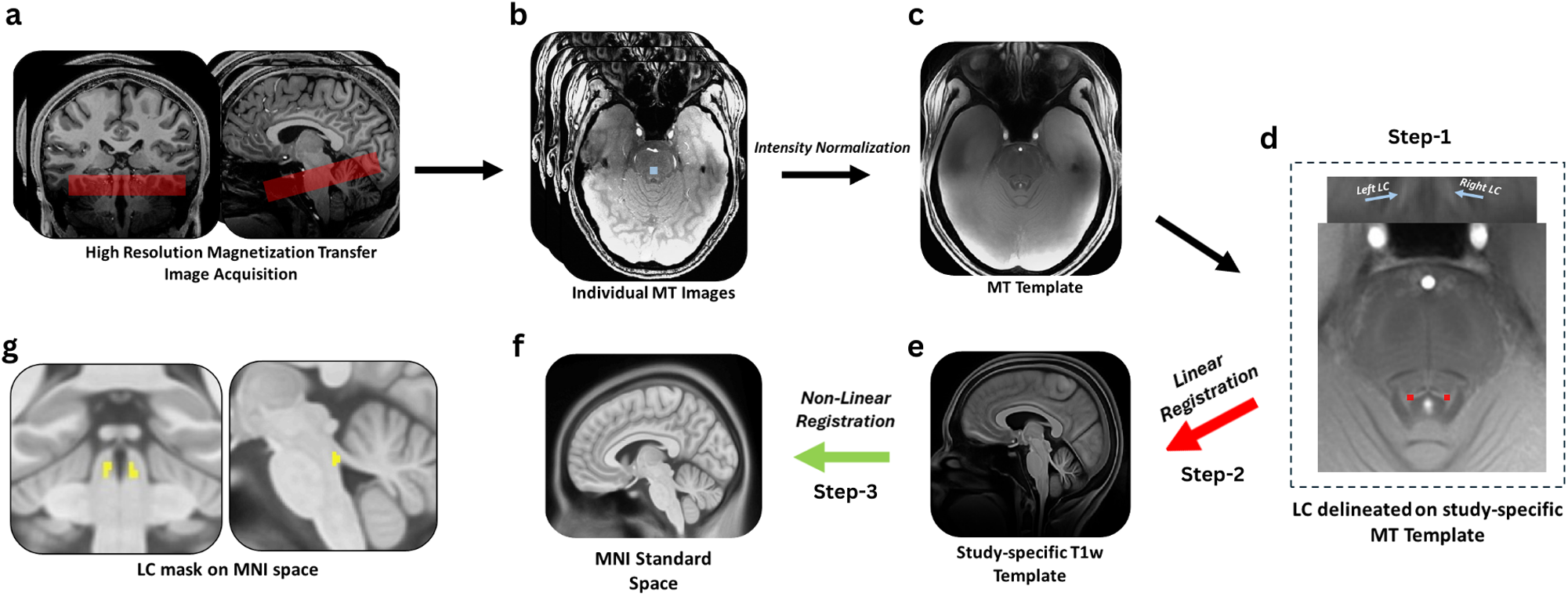
Locus Coeruleus delineation pipeline. MT-TFL pre-processing and spatial transformation pipeline. (a) High resolution MT-TFL images acquired by positioning the approximately perpendicular to the pons, encompassing the region between the inferior colliculus and the inferior border of the pons. Panel (b) shows an example of individual MT-TFL images acquired using this sequence with 10×10 voxel (blue) placed on pons. Panel (c) demonstrates a study-specific MT-TFL template created after intensity normalizing the individual MT-TFL images. Panel (d) shows left, and right LC delineated on subject-specific MT-TFL template using ITK-SNAP. Panel (e) depicts study-specific T1w template created to act as a bridge to transform LC mask to MNI space, red arrow indicates MT template and was rigidly registered to the whole-brain study-specific T1w template. Panel (f) shows MNI space where T1w template was non-linearly registered (green arrow). Panel (g) depicts left and right LC mask transformed to MNI space after using the transformation matrices obtained from step-2 and step-3.

#### Spatial transformation of LC

To facilitate spatial transformations across scans acquired at different resolutions, LC-sensitive structural images and LC segmentations were resampled to match the isotropic voxel resolution of the T1w images using the *mri_convert* function in FreeSurfer (Version 7.1; https://surfer.nmr.mgh.harvard.edu/, Martinos Center for Biomedical Imaging, Charlestown, Massachusetts). This resampling minimized interpolation errors and ensured accurate alignment during subsequent registration steps.

All spatial transformations were conducted using the deformable SyN algorithm in ANTs (Klein et al., 2009), known for its high precision and flexibility. First, the study-specific MT-TFL images were linearly registered to each subject’s native-space whole-brain T1-weighted images using a rigid transformation via the antsRegistrationSyN function (Step 2 – red arrow). As MT-TFL and T1w images were acquired in the same session and from same individuals, a rigid registration (following Yi et al., 2023) was sufficient to ensure accurate anatomical alignment while minimizing distortion. The T1w template served as an intermediate bridge to facilitate the transformation of MT-TFL images into MNI space. Next, the whole-brain study-specific T1w template (Figure 2e) was non-linearly registered to MNI space (Figure 2f) using the *antsRegistrationSyN* function (Step 3 – green arrow) to account for individual anatomical variability and ensure proper standardization to the MNI152 standard space (Tustison et al., 2010; Avants et al., 2011). The transformation matrices obtained from step 2 and 3 during registration steps (Figure 2e and Figure 2f) were then applied to transform the LC segmentations, delineated on MT-TFL LC-sensitive structural scans into MNI space using *antsApplyTransforms* in a single step (Figure 2g). Individual LC segmentations were transformed using the nearest neighbor option to avoid creating ambiguous voxels during the interpolation process (Yi et al., 2023), ensuring clear association with the original LC segmentation.

#### fMRI analyses

Analyses of fMRI data during preprocessing and statistical modeling steps were conducted using fMRIPrep version 22.0.2 (Esteban et al. 2019) and Statistical Parametric Mapping 12 (SPM12; Wellcome Centre for Human Neuroimaging, University College, London, UK, 2012), respectively. The raw DICOM data were converted to NIfTI format, preserving the original imaging parameters. The initial three volumes were discarded to eliminate transient effects in the MR signal. Preprocessing of the functional data was performed using the default fMRIPrep pipeline (Esteban et al. 2019), which includes slice timing correction, realignment, co-registration with the T1w image using boundary-based registration (FreeSurfer; Greve and Fischl 2009), normalization to MNI standard space. Due to small size of the LC, smoothing was performed with a 2mm FWHM Gaussian kernel, following the recommendation to use a kernel of at least 1.5 times the native voxel size (Worsley and Friston, 1995; Yi et al., 2023). The choice of using a 2mm smoothing kernel was to ensure maintaining spatial precision to avoid signal blurring (Yi et al., 2023).

The first-level analysis was performed using SPM12 in Matlab (MathWorks). For each participant, contrast images were modeled in the framework of the general linear model (GLM) for each of the seven conditions, ranging from 100% fear to 100% happiness, while ambiguity levels varied from 30% fearful – 70% happy to 70% fearful – 30% happy in 10% increments, including a balanced 50% fearful – 50% happy condition. Each event was modeled based on the onset of face trials and convolved with the canonical hemodynamic response function (HRF). To account for potential confounding factors, head motion parameters (6 motion regressors) were included as nuisance regressors, minimizing the influence of motion-related artifacts. Additionally, first five aCompCor regressors were used to account for physiological noise in the data (Behzadi et al., 2007), since physiological measurements (e.g., cardiac and respiratory signals) were not available. To minimize the impact of movement artifacts, which high-resolution functional images are particularly susceptible to (Havsteen et al., 2017), we excluded subjects with severe motion (mean framewise displacement ≥ 0.5mm, Power et al., 2011). This conservative approach allowed for a more accurate modeling of neural activity associated with each condition (Siegele et al., 2013).

First, the main effects of each condition were examined to identify the whole brain activation. Group data were analyzed using a random-effects model to examine activation patterns across conditions. Parametric *t*-maps were generated to identify brain regions significantly activated for each condition relative to baseline in SPM. One-sample *t*-tests were used to analyze group-level effects across age for absolute ambiguity condition, while two-sample *t*-tests compared neural responses between young and older adults separately for each condition. Additionally, age-related differences in neural responses were also calculated by grouping conditions to three conditions: anchor condition (100% fearful and 100% happy), intermediate ambiguity condition (30%, 40%, 60% and 70%) and absolute ambiguity condition (50% fearful – 50% happy). For the whole-brain exploratory analysis, voxel-wise statistical maps were initially thresholded at *p* < 0.001 (uncorrected) as the cluster-defining threshold (CDT). Multiple comparisons were corrected using the false discovery rate (FDR) at *p* < 0.05, which controls the expected proportion of false positives among significant results and increases sensitivity to true activation clusters (Genovese et al., 2002; Chumbley et al., 2009). All analyses were performed with gender and years of education as a covariate and a gray matter mask was used for whole-brain analyses (Fonov et al., 2011). Cortical and subcortical regions were defined using anatomical masks extracted from the AAL3 atlas (Rolls et al., 2020) and all results are reported in MNI space.

For the target analysis of the LC, we applied an inclusive brainstem mask (Beissner et al., 2013). Given the small size of the LC, we applied small volume correction (SVC) to evaluate significant activations from the study-specific LC mask in the second-level analyses (Figure 2g). Voxel-wise statistical maps were initially thresholded at *p* < 0.005 (uncorrected) as the CDT. To correct multiple comparisons, significant clusters were identified using family-wise error (FWE) correction at *p* < 0.05, ensuring stringent control for false positives, particularly in hypothesis-driven analyses (Nichols & Hayasaka, 2003).

#### Functional connectivity analyses

To further explore the connection between the LC and prefrontal areas, we conducted functional connectivity (FC) analyses. The regions of PFC such as dlPFC and dmPFC were selected as regions of interest due to their well-established roles in conflict resolution (Grueschow et al., 2020; Sun et al., 2023). Additionally, we performed control analyses with other ROIs, including the ventromedial prefrontal cortex (vmPFC), insula, anterior cingulate cortex (ACC) given their involvement in conflict processing, processing emotional ambiguity and decision-making (Sun et al., 2017; Suzuki & Tanaka, 2021; Wu et al., 2021; Ince et al., 2023), and the fusiform face area (FFA), given its involvement in facial perception (Lee et al., 2014; Lee et al., 2018).

We employed Generalized Psychophysiological Interaction (gPPI) analysis using the CONN toolbox (v22.a, Nieto-Castanon & Whitfield-Gabrieli 2022). These analyses were guided by our initial hypothesis regarding age-related changes in the LC connectivity with broader cortical PFC regions. The gPPI approach examines condition-specific changes in connectivity by modeling the interaction between psychological variables (e.g., task conditions) and the physiological time series extracted from the seed region. Given the evidence of lateralization in LC connectivity linked to cognitive performance, emotional regulation, and neurodegenerative conditions (Veréb et al., 2023; Jacobs et al., 2015; Sun et al., 2023) and our results indicating greater activity within the left LC in older compared to younger adults during absolute ambiguity condition—we analyzed the left and right LC as separate seed regions. Target ROIs included the left and right dlPFC, dmPFC, vmPFC, insula, ACC, and FFA. The connectivity profiles of the left and right LC (seeds) were assessed using the corresponding time series from the left and right hemispheres separately to the target ROIs across each of the seven emotional conditions grouped into three conditions as per behavioral and whole-brain analyses. The anchor (unambiguous condition included 100% happy, 100% fearful; intermediate ambiguity condition included 30%-70% and 40%-60% happy-fearful expressions in either direction, and absolute ambiguity included 50% happy - 50% fearful conditions). The gPPI model included all conditions, generating beta coefficients that reflect the strength of connectivity between the seed and target ROIs under each condition. Additionally, gPPI measures represent the level of task-modulated connectivity between two ROIs (i.e., changes in functional association strength covarying with the external or experimental factor). gPPI is computed using a separate multiple regression model for each target ROI timeseries (outcome). Each model includes as predictors: a) all the selected task effects (main psychological factor in PPI nomenclature); b) each seed ROI-BOLD timeseries (main physiological factor in PPI nomenclature); and c) the interaction term specified as the product of (a) and (b) (PPI term; Nieto-Castanon, 2020). Results were initially thresholded using a combination of a connection and cluster-forming threshold at *p*<0.05. Significant clusters were identified and corrected for multiple comparisons using cluster-level FDR correction at p < 0.05 (Chumbley et al., 2009).

#### Statistical Analysis

LC activity during the absolute-ambiguity condition was correlated with both task-dependent RTs and task-independent emotional well-being measures, within and across younger and older adults. Partial correlations, controlling age, gender, and education years, were computed using pingouin.partial_corr in Python. Additionally, we examined the functional connectivity of the left and right LC separately with left and right target regions (dlPFC, dmPFC, vmPFC, insula, ACC, and FFA) and assessed their relationship with task-independent mental well-being and emotional resilience index using partial correlations, controlling for the same covariates. Group differences in these correlations were tested using the Chow test (Chow, 1960) to evaluate differences in regression coefficients between younger and older adults. To enhance the robustness of our results (Davison & Hinkley, 1997), both correlation estimates, and regression model comparisons were bootstrapped with 1,000 iterations using scipy.stats.bootstrap in Python to obtain confidence intervals. Outliers were assessed using the Interquartile Range (IQR) method, and all data points were found to fall within the range of −3 to +3, indicating no outliers.

## Results

### Behavioral

#### Age-Related performance during emotional ambiguity processing

Across all participants, ambiguous faces were perceived as more fearful than happy (*xhalf* = 51.16 ± 6.68). However, two-sample *t*-test analyses revealed that older adults exhibited significantly lower xhalf values (xhalf = 49.45 ± 6.03) compared to younger counterparts (*xhalf* = 53.08 ± 6.05, Figure 1c), indicating that older adults were more inclined to perceive ambiguous faces as happy than fearful (*t*_(142)_ = −3.06, *p* = 0.0004, *d* = −0.60). In contrast, the steepness of the psychometric curve (α) did not significantly differ between age groups (*t*(142) = 1.33, *p* = 0.186, *d* = 0.21), suggesting that while older adults had a bias toward perceiving ambiguous faces as happy, their overall sensitivity to changes in emotional intensity remained comparable to younger adults.

A 2 (age group: young and old) by 3 (ambiguity level: anchor, intermediate, and absolute-ambiguous) repeated-measures analysis of variance (ANOVA) was conducted with RT as the dependent variable. The analysis revealed no significant main effect of age group, *F*_(1, 142)_ = 0.31, *p* = 0.57, η² = 0.002, but a significant main effect of ambiguity level, *F*_(2, 284)_ = 79.36, *p* = 4.15 × 10C^28^, η² = 0.5 (Figure 1f). Follow-up analyses revealed that RTs were significantly longer for absolute-ambiguous faces compared to intermediate ambiguous (*t*_(142)_ = 7.08, *p* = 1.41 × 10^11^, Hedge’s g = 0.39) and anchor faces (*t_(_*_142)_ = 9.95, *p* = 1.11 × 10^-18^, Hedge’s g = 0.96) regardless of age. Additionally, no significant interaction between age group and conditions were observed, *F*_(2, 284)_ = 0.33, *p* = 0.71, η² = 0.002, indicating that the effect of different ambiguity levels on RT was similar across younger and older adults.

A similar ANOVA, with confidence ratings as the dependent variable, revealed a significant main effect for age group, *F*_(1, 142)_ = 10.48, *p* = 0.0015, η² = 0.07 with post hoc analysis suggesting lower confidence in older compared to younger adults regardless of ambiguity condition (*t*_(142)=_ −11.84, *p* = 1.02 × 10^-11^, Hedge’s *g = −0.17*). Additionally, significant main effect of ambiguity level was also observed *F*(2, 284) = 441.78, *p* = 1.26 × 10^-87^, η² = 0.75 with post hoc analyses indicating that absolute-ambiguity resulted in significantly lower confidence compared to unambiguous conditions (*t*_(142)_ = 40.88, *p* = 4.01 × 10^-12^, Hedge’s *g* = 1.09) and intermediate ambiguity condition (*t*_(142)_ = 15.37, *p* = 4.01 × 10^-12^, Hedge’s *g* = 0.32) regardless of age. The interaction between age group and ambiguity level did not reach significance, *F*_(2, 284)_ = 2.36, *p* = 0.097, η² = 0.16, indicating that the effect of ambiguity levels on confidence was similar for both younger and older adults.

### Age-related improvement in emotional well-being

Our analysis of emotional well-being revealed that older adults had significantly better ‘Emotional Resilience Index’ derived from the PCA analysis (*t*_(142)_ = −8.32, *p* < 0.001, *d* = 1.40), indicating greater emotional well-being compared to younger adults. In contrast, PCA of Cognitive Function Index showed that younger, compared to older, adults performed better in cognitive tasks (*t*_(142)_ = 6.382, *p* < 0.001, *d* = 1.07). All individual measurements and differences between two age groups are reported in <u>Table S1</u>.

#### fMRI results

### Neural underpinnings of emotional ambiguity across age

First, we examined emotional ambiguity processing across age groups using a one-sample *t*-test. The results revealed significant activation in several key regions, including the bilateral middle frontal gyrus, inferior frontal gyrus, superior frontal gyrus, insula, superior temporal gyrus, middle temporal gyrus, postcentral gyrus, precentral gyrus, angular gyrus, and putamen (<u>Figure S1</u>). Additionally, we observed significant activation in the bilateral dorsolateral prefrontal cortex (dlPFC), which is marked in <u>Figure S1</u>, highlighting its role in emotional ambiguity processing.

### Age-related differences during emotional absolute-ambiguity processing

Next, to examine our primary hypothesis on the age-related differences during ambiguity processing, a two-tailed *t*-test was conducted. Younger, compared to older adults, exhibited significantly greater activation in regions associated with sensory processing, cognitive control, and emotional regulation, including the left postcentral gyrus, bilateral insula, bilateral middle frontal gyrus, bilateral inferior frontal gyrus, right dorsolateral superior frontal gyrus, right angular gyrus, left hippocampus, right caudate, left precentral gyrus, and right superior medial gyrus (Table 1a). In contrast, older adults demonstrated greater activation in the right fusiform gyrus, right lingual gyrus, and left thalamus, compared to younger adults (Table 1b) (<u>Figure S4</u>).

**Table 1:**
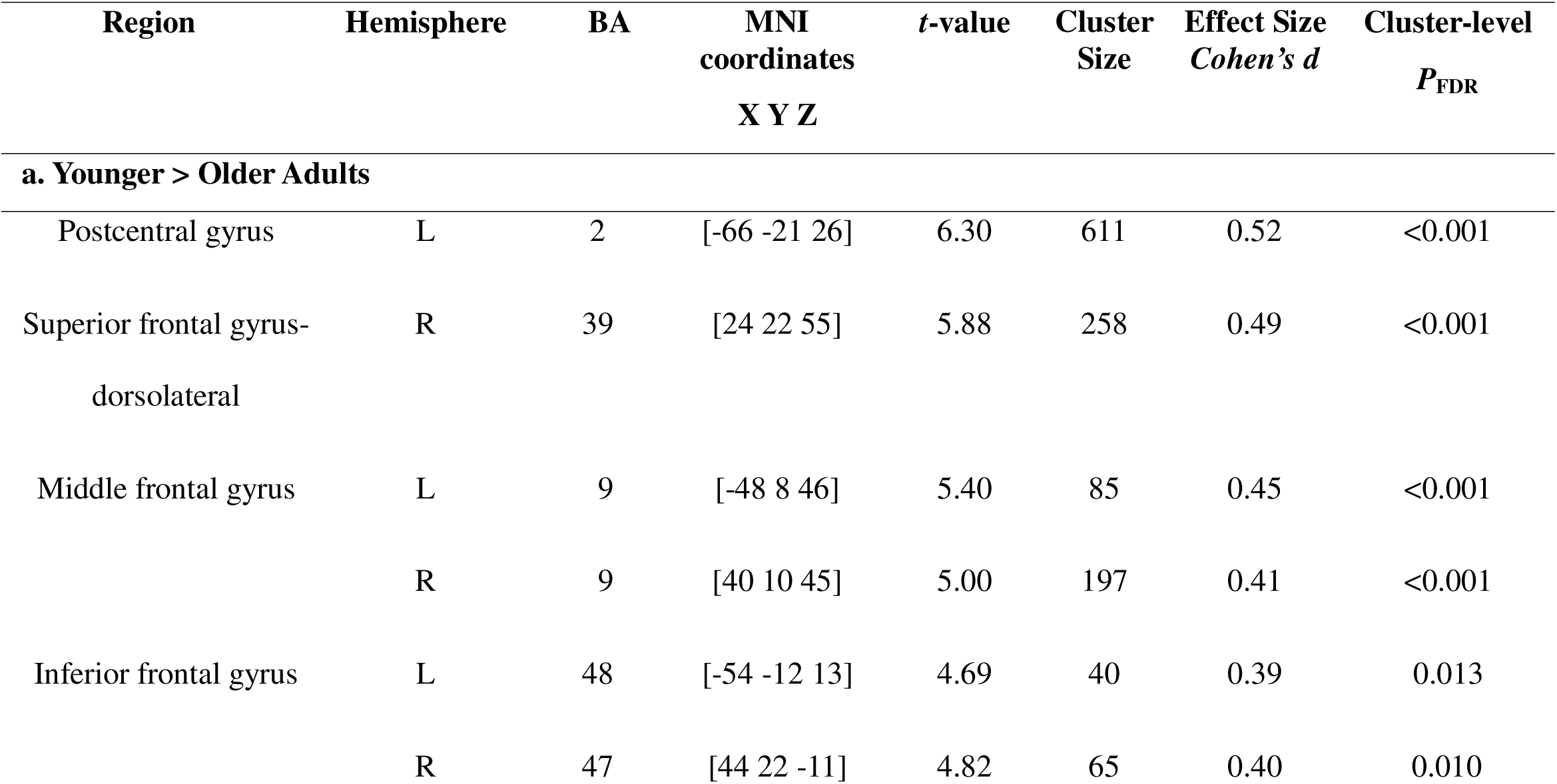
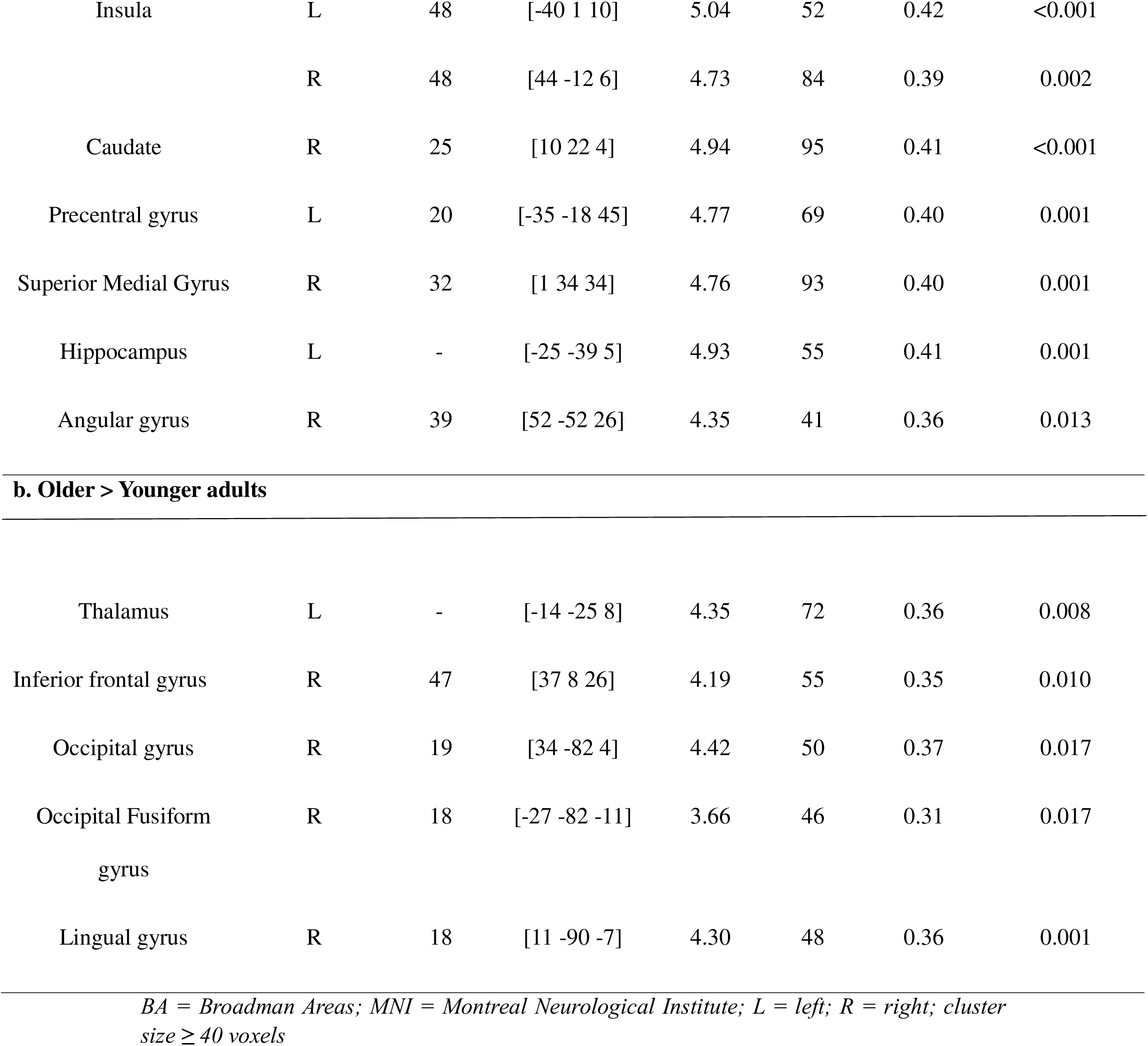
Peak coordinates for significant clusters from whole brain analyses during absolute ambiguity condition.

Although not the primary focus of this study, we examined age-related differences in brain activation during the anchor and intermediate conditions. For the anchor condition, younger adults exhibited heightened recruitment in regions such as the bilateral insula and left postcentral gyrus compared to older adults (Table 2a). Conversely, older adults demonstrated greater activation in posterior and visual regions, including the bilateral fusiform gyrus, posterior cingulate cortex, bilateral calcarine cortex, and bilateral precentral gyrus compared to younger adults (Table 2b). For the intermediate condition, younger adults showed increased engagement in areas such as the left postcentral gyrus, mid cingulate cortex, bilateral insula, bilateral supramarginal gyrus, left hippocampus, and right middle frontal gyrus compared to older adults (Table 3a). In contrast, older adults exhibited greater activation in regions such as the bilateral caudate, thalamus, and occipital fusiform gyrus compared to younger adults (Table 3b). In none of these analyses LC was significantly activated.

**Table 2:**
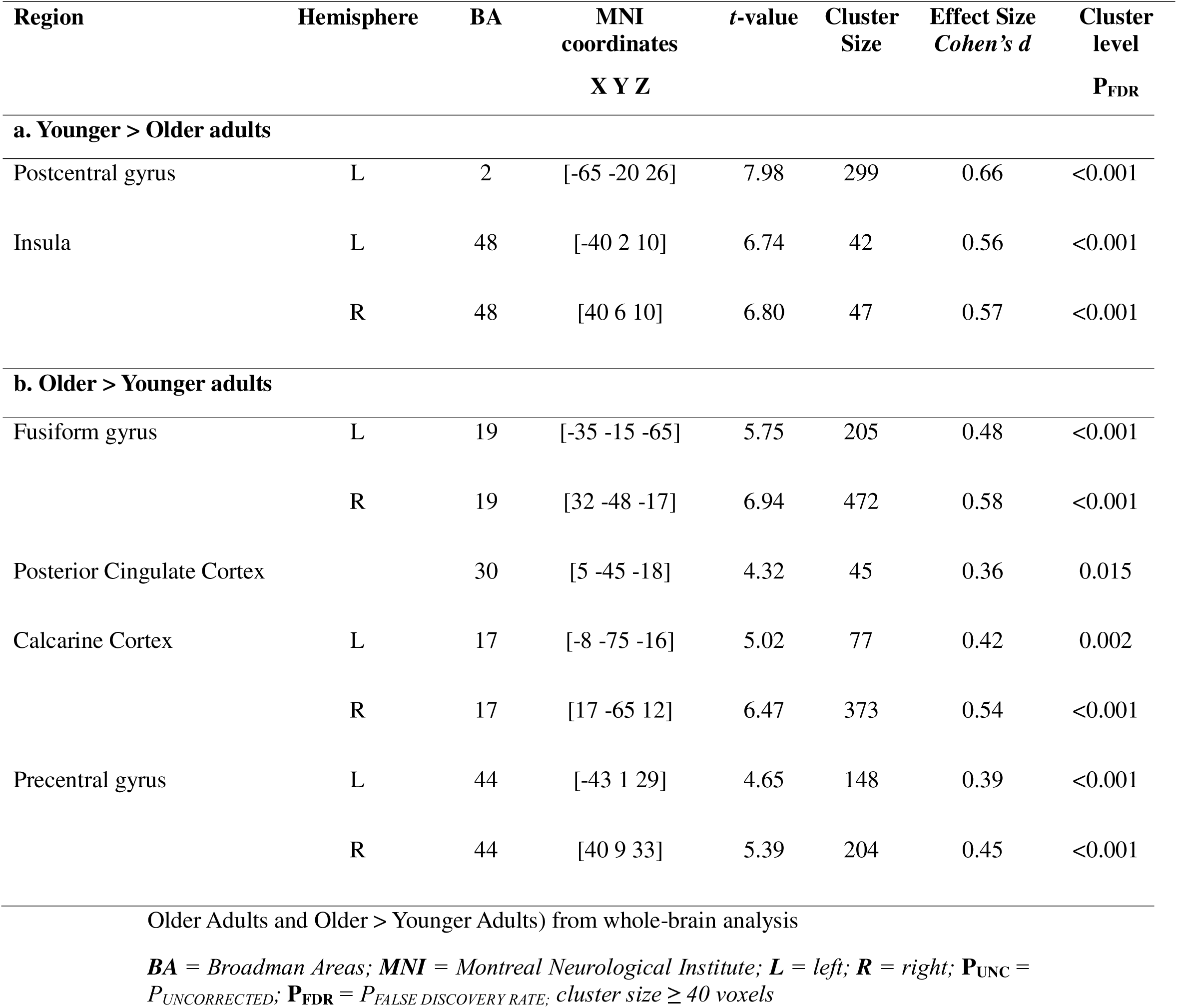
Peak coordinates of significant clusters for the anchor condition contrast (Younger >.

**Table 3:**
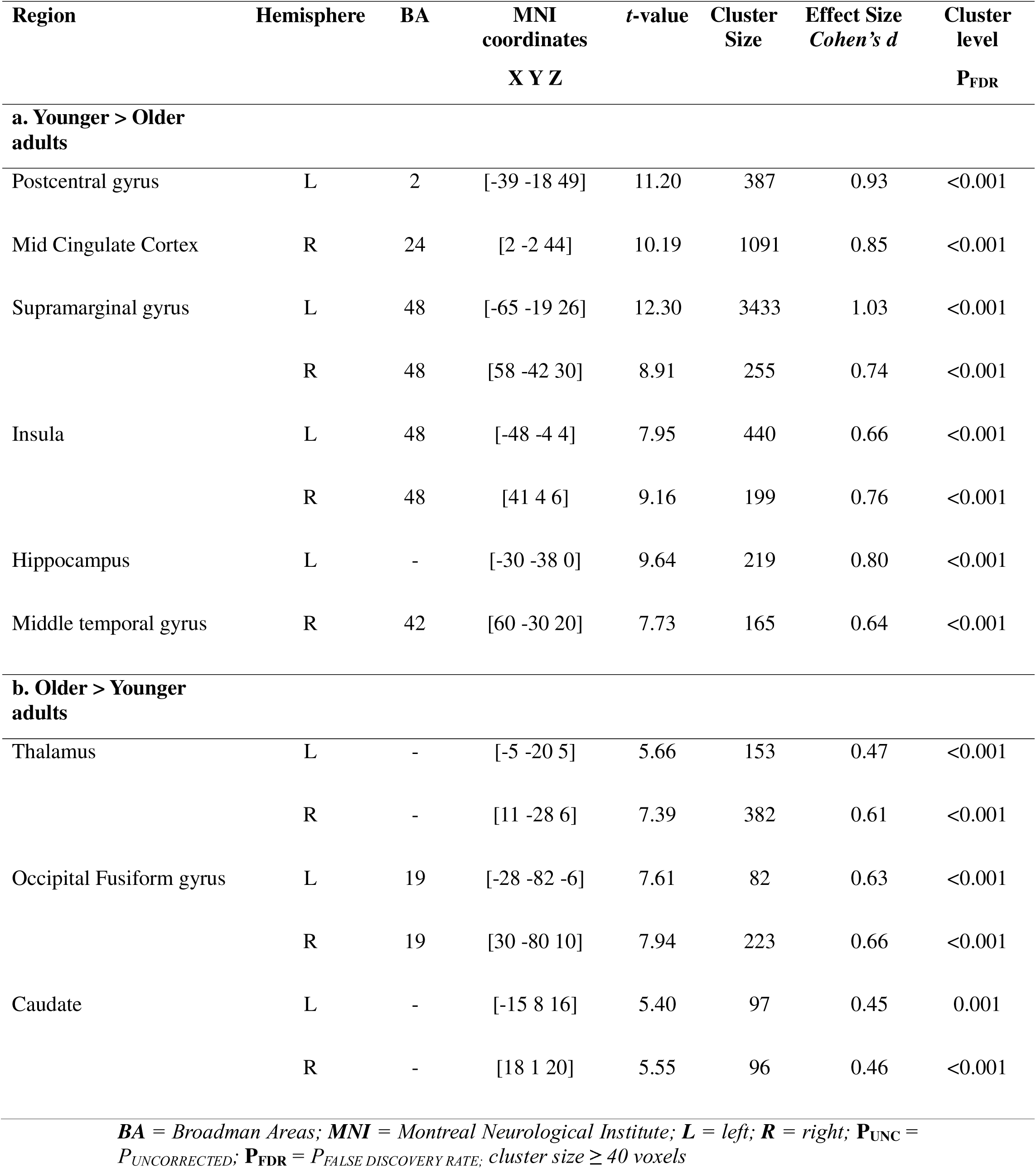
Peak coordinates of significant clusters for intermediate ambiguity condition contrast: younger > older adults and older > younger adults (whole brain analysis)

### Higher LC activation in older compared to younger adults during emotional absolute-ambiguity processing

In line with our second hypothesis, during the absolute-ambiguity condition, we observed significant higher activity in the left LC among older adults compared to younger adults [Figure 3a; *t* = 3.51, *P_FWE_ = 0.024, cluster size = 6,* MNI-coordinates of peak voxel, x = −5; y = −38; z= −24], with SVC confirming that 75% of the significant activated voxels was localized within the study-specific LC mask, while the remaining 25% extended outside the mask (see <u>Figure S2</u>). Full pattern of brainstem activation can be seen in Figure 4 (Table 4). Additionally, it has to be noted that when examining brainstem activity across all participants, we did not observe LC activation, suggesting individual variability in LC responsivity may mask group-level effects (Table S2). Importantly, we also did not find any age-related significant differences in the LC activity in either the emotionally arousing fearful and happy (anchor) or the intermediate ambiguity conditions, all *p*s > 0.05.

**Figure 3:**
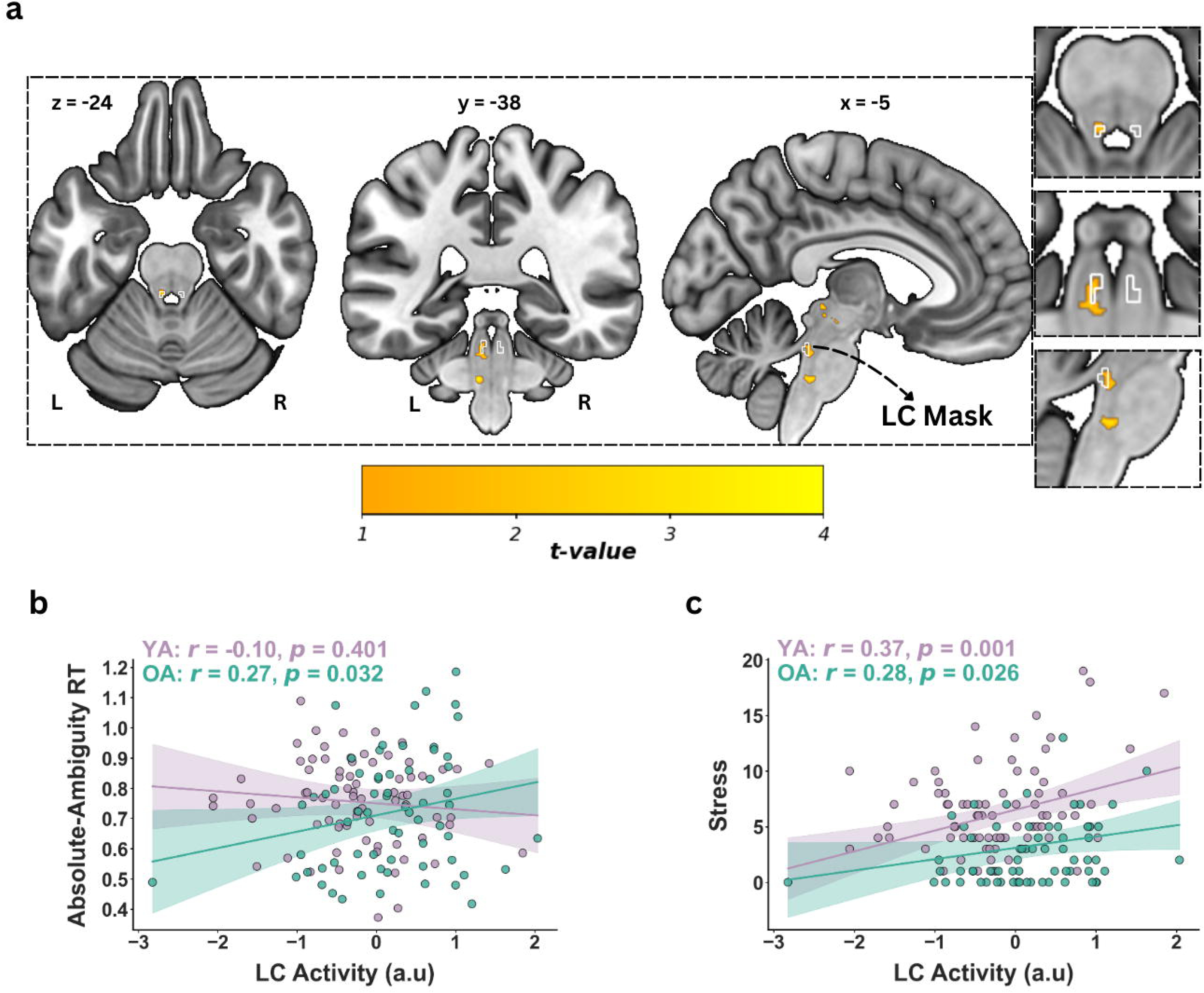
LC activation and behavioral correlation. (a) Higher Locus Coeruleus (LC) activation in older (OA) > younger adults (YA) during absolute-ambiguity condition. Significant activation thresholded with *P_FWE_* <0.05 within LC mask (white). Panel (b) illustrates LC activity and its link with response time (RT) during absolute-ambiguity condition for both YA and OA. Panel (c) shows LC activity and its link with self-reported stress measures in YA and OA. FWE = family wise error.

**Figure 4:**
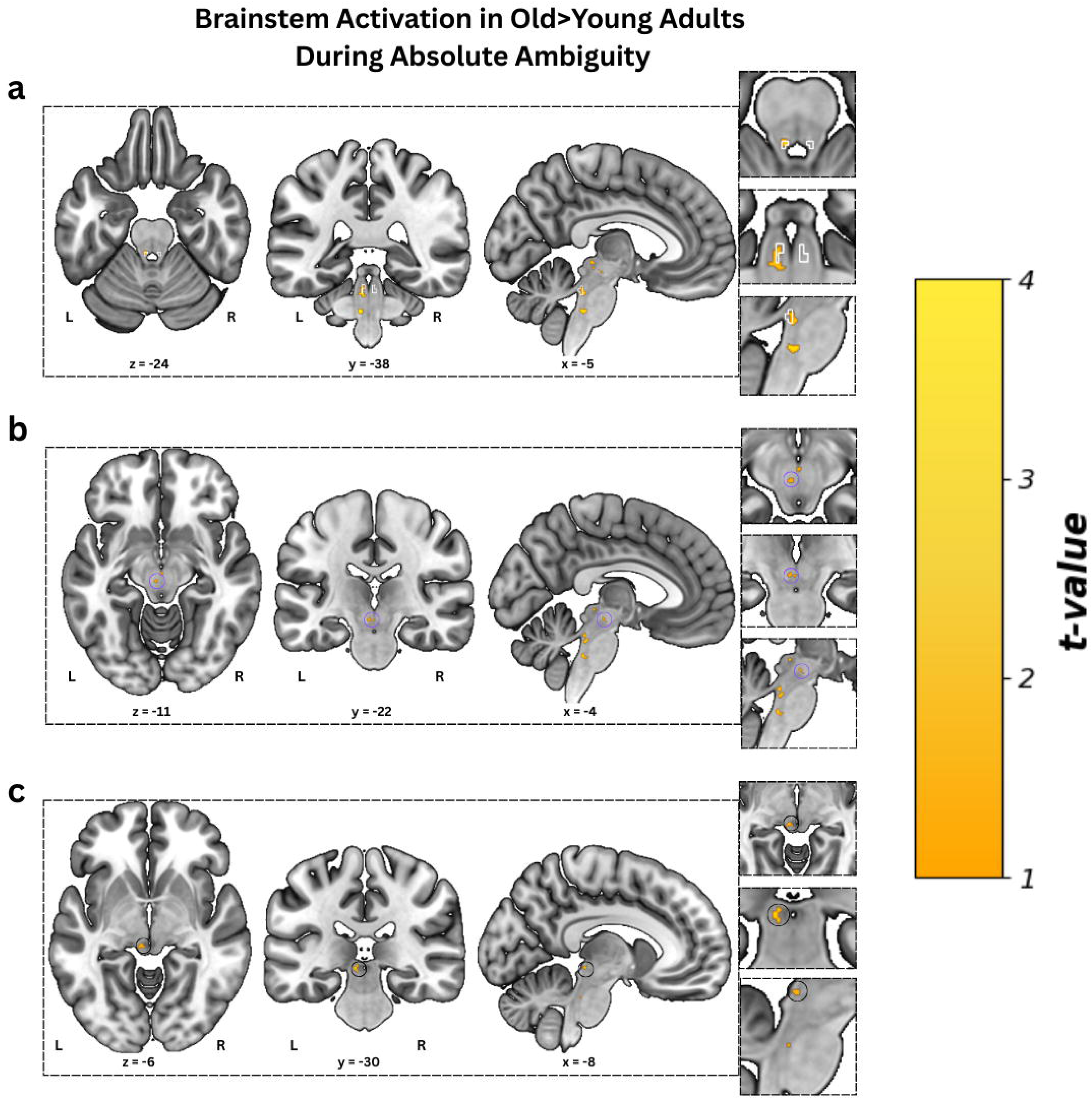
Brainstem Activation in Old>Young Adults During Absolute Ambiguity Condition. Whole-brainstem activation map for the Old > Young contrast during the absolute ambiguity condition, thresholded at *p < 0.005* uncorrected with a voxel extent threshold of ≥10 voxels. Panel (a) Activation cluster overlapping with the locus coeruleus within LC mask (white). Panel (b) Cluster observed near the left red nucleus (purple circled). Panel (c) Cluster observed near the left midbrain (black circled).

**Table 4:**
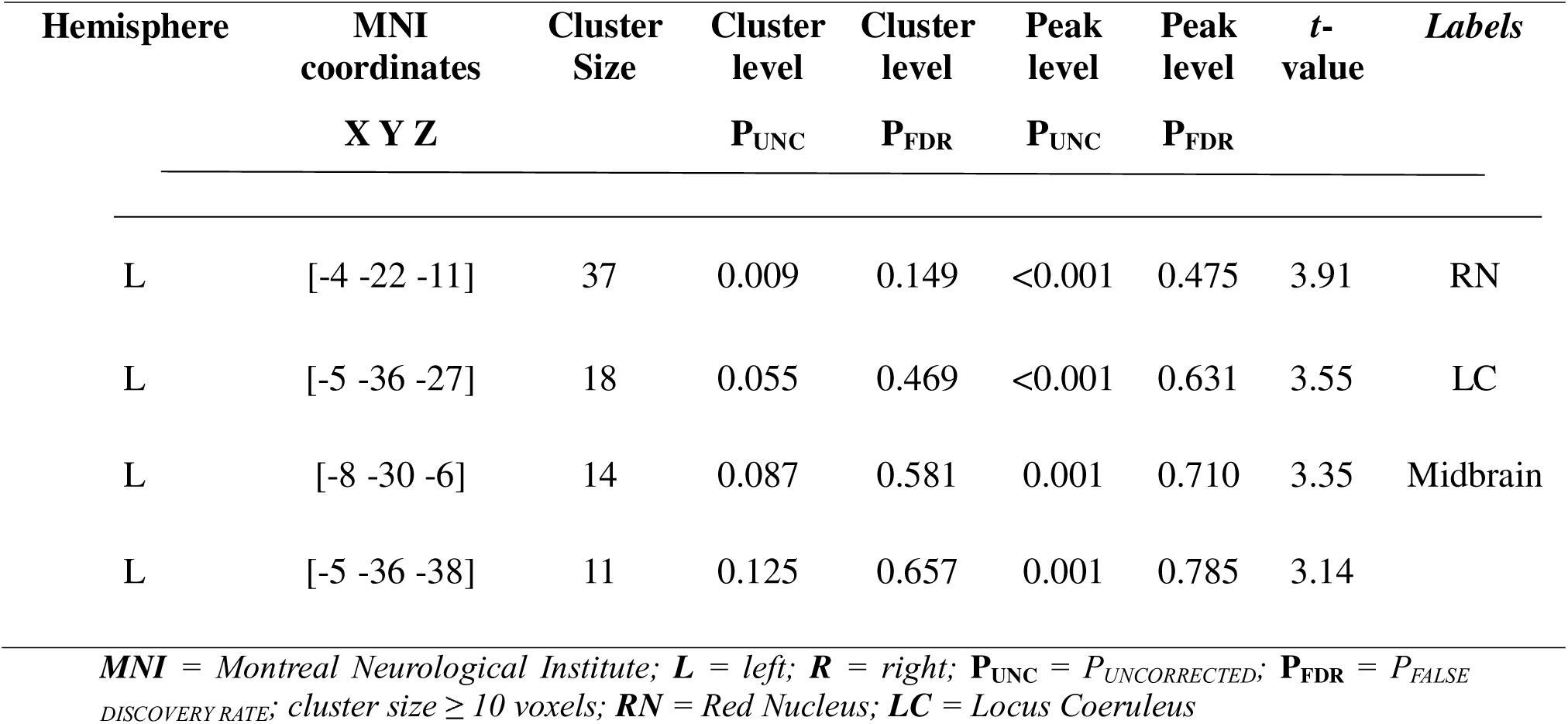
Peak coordinates of activated clusters in the brainstem for absolute ambiguity condition in Old>Young contrast.

### Association between task-dependent RTs and LC activity

First, we investigated the relationship between left LC activity and task dependent RTs during absolute-ambiguity condition, both across the entire cohort and separately for younger and older adults. Across the entire group, we found no significant correlation between the left LC activity and RTs during absolute-ambiguity condition (*r* = 0.08, *p* = 0.37, 95% CI: [-0.09, 0.24]). To determine whether the relationship between the left LC activity and RTs during absolute-ambiguity processing differed significantly between age groups, we compared regression coefficients between younger and older adults. Bootstrapped analysis confirmed a significant difference (*F*_(5,134)_ = 3.99, *p* = 0.004, 95% CI: [1.09, 8.63]), suggesting a stronger relationship between left LC activity and RTs in older adults. Within age group analyses revealed that older adults showed positive correlation between the left LC activity and RTs during absolute-ambiguity processing (Figure 3b; *r* = 0.27, *p* = 0.032, 95% CI: [0.02, 0.49]). A bootstrapped correlation analysis using 1,000 iterations confirmed the robustness of these findings, showing consistent results (mean *r = 0.278, 95% CI*: [0.018, 0.504]) in older adults. This indicates that, for older adults, higher left LC activation was associated with longer RTs during the absolute-ambiguity condition. In contrast, within younger adults, the relationship was not significant (*r* = −0.10, *p* = 0.401, 95% CI: [-0.33, 0.14]).

### Association between emotional well-being and LC activity

Next, we correlated the left LC activity with task independent emotional well-being measures. Across the entire group, we observed a significant positive correlation between left LC activity and task independent self-reported stress scores (*r* = 0.33, *p =* < 0.001, 95% CI: [0.17, 0.47]). This suggests that, overall, higher left LC activity was associated with greater self-reported stress levels across the cohort regardless of age. Within-group analyses also revealed significant positive correlations between stress scores and left LC activity in both younger (*r* = 0.37, *p* = 0.001, 95% CI = [0.15, 0.55]) and older adults (*r* = 0.28, *p* = 0.026, 95% CI: = [0.04, 0.50]; see Figure 3c). Additionally, our analysis confirmed a significant difference in regression coefficients between the left LC activity and stress scores between age groups (*F*_(5,134)_ = 6.80, *p =* < 0.001, 95% CI: [2.67, 12.69]), confirming that the association between LC activity and task independent stress scores was more pronounced in younger adults.

### Higher LC-dlPFC in older compared to younger adults during emotional absolute-ambiguity processing

To address our third hypothesis and further investigate age-related differences in the LC connectivity with PFC regions, we compared left and right LC connectivity separately with left and right dlPFC, dmPFC, vmPFC, insula, ACC or FFA between younger and older adults during the absolute-ambiguity condition.

Results revealed a significant age-related increase in connectivity between the left LC and right dlPFC, with older adults exhibiting stronger connectivity compared to younger adults (Figure 5). β = 0.12, *t* = 2.26, *p* = 0.024, *P_FDR_*= 0.045). However, no significant age-related differences were found for the left LC and left dlPFC connectivity (β = 0.09, *t* = 1.40, *p* = 0.16) or left LC connectivity with the bilateral dmPFC (right: β = 0.10, *t* = 1.40, *p* = 0.16; left: β = 0.11, *t* = 1.11, *p* = 0.27). Similarly, no significant age-related differences were found for the LC connectivity with other areas such as the vmPFC, insula, ACC or FFA (all *p*s > 0.05) in absolute-ambiguity condition.

**Figure 5:**
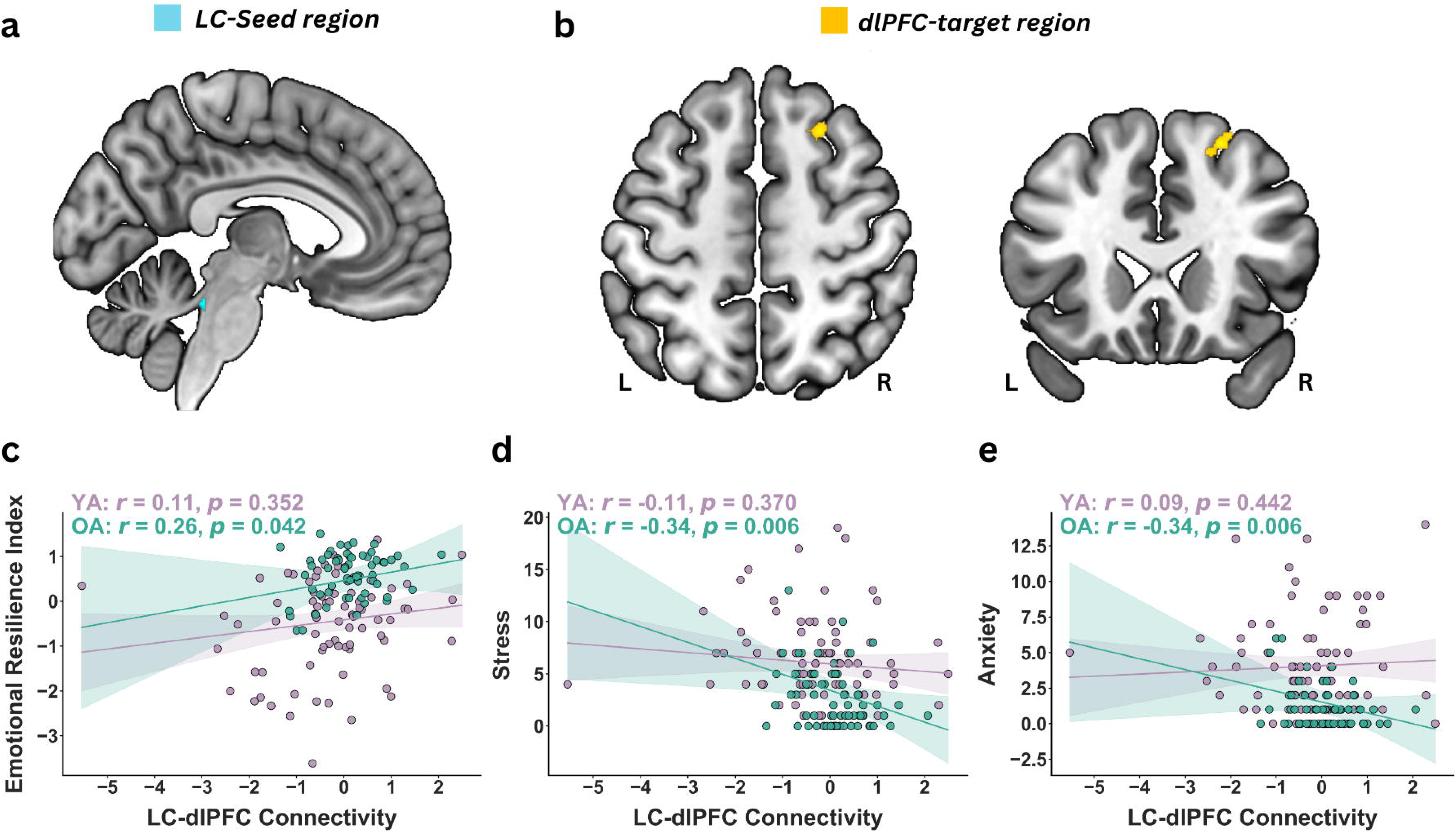
LC-dlPFC connectivity and behavioral correlation. Absolute-ambiguity elicits greater LC-dlPFC connectivity in older adults (OA) compared to younger adults (YA). Panel (a) shows the Locus coeruleus (LC) seed-region shows greater connectivity with panel (b) showing the right dorsolateral prefrontal cortex (dlPFC) target regions in ROI-to-ROI analysis using generalized psychophysiological interaction. Panel c to e illustrates the relationship between LC-dlPFC connectivity and mental well-being measures in OA: Panel (c) depicts that higher LC-dlPFC connectivity is positively associated with greater emotional resilience, while panel (d) shows that it is negatively associated with stress, and panel (e) with anxiety scores. These significant associations are observed in OA, but not in YA.

Additionally, no significant age-related differences were found for right LC connectivity with any target regions, including bilateral dlPFC, dmPFC, vmPFC, insula, ACC and FFA (all p > 0.05), suggesting that the observed age-related effect on LC-dlPFC connectivity was specific to the left LC–right dlPFC pathway. Also, across age groups, no significant functional connectivity was observed between left or right LC with any of the target seeds (dlPFC, dmPFC, vmPFC, insula, ACC or FFA) during any of the conditions; anchor, intermediate or absolute ambiguity conditions (all *p*s > 0.05).

While a significant age-related difference in functional connectivity between the left LC and right dlPFC was observed at a CDT of *p* < 0.05, it did not survive stricter thresholds (*p* < 0.005 or *p* < 0.001), indicating the need for cautious interpretation due to the potential for inflated false positives (Eklund et al., 2016).

### Higher LC-dlPFC connectivity links with better mental well-being and emotional resilience among older adults

Finally, we examined whether LC-dlPFC connectivity during the absolute-ambiguity condition was associated with task independent mental well-being scores and emotional resilience index, both across the entire group and separately within younger and older adults’ groups.

### LC-dlPFC connectivity and task-independent stress scores

Across all participants, stronger left LC–right dlPFC connectivity showed a trend-level negative association with lower self-reported stress scores (*r* = −0.15, *p* = 0.086, 95% CI: [-0.31, 0.002]). Within-group analyses revealed that this relationship was significant among older adults (Figure 5d; *r* = −0.34, *p* = 0.006, 95% CI: [-0.54, −0.10]) whereas no association was observed in younger adults (*r* = −0.11, *p* = 0.37, 95% CI: [-0.33, 0.12]). Additionally, bootstrapped analysis confirmed a significant difference in regression coefficients between age groups (*F*_(5,134)_ = 4.59, *p* = 0.003, 95% CI: [1.34, 9.16]), indicating that age-related increase in left LC - right dlPFC connectivity links with reduced stress scores.

### LC-dlPFC connectivity and task-independent anxiety scores

Across all participants, left LC – right dlPFC connectivity was not significantly associated with self-reported anxiety scores (*r* = −0.01, *p* = 0.90, 95% CI: [-0.16, 0.18]). However, within-group analyses revealed an age-specific effect. Among older adults, stronger left LC– right dlPFC connectivity was significantly associated with lower anxiety scores (*r* = −0.34, *p* = 0.006, 95% CI: [-0.54, −0.11]). In contrast, no such association was found in younger adults (*r* = 0.09, *p* = 0.442, 95% CI: [-0.14, 0.31]). Bootstrapped analysis revealed a significant difference in regression coefficients (*F*_(5,134)_ = 7.76, *p = < 0.001,* 95% CI: [3.30, 14.56]), confirming that the association between left LC connectivity with right dlPFC connectivity and self-reported anxiety scores differed significantly between age groups, suggesting that age-related increase in left LC - right dlPFC connectivity links with reduced anxiety scores.

### LC-dlPFC connectivity and emotional resilience index scores

Across all participants, stronger left LC–right dlPFC connectivity showed a trend-level positive association with a greater task-independent emotional resilience index (*r* = 0.16, *p* = 0.063, 95% CI: [-0.010, 0.32]). Within-group analyses revealed that this relationship was significant in older adults (*r* = 0.26, *p* = 0.042, 95% CI: [0.01, 0.47]), whereas no significant association was found in younger adults (*r* = −0.11, *p* = 0.355, 95% CI: [-0.33, 0.12]; Figure 5c). This was confirmed by the difference in regression coefficients between age groups using bootstrapped analysis (*F*_(5,134)_ = 7.75, p = < 0.001, 95% CI: [3.61, 14.20]), indicating that the relationship between left LC–right dlPFC connectivity and the emotional resilience index significantly differs between age groups. Further suggesting that age-related increases in left LC – right dlPFC connectivity contribute to better emotional resilience in older adults.

Additional analyses were conducted using the cognitive function index, which incorporates cognitive and executive control measures (reported in Table S1). Across all participants, left LC – right dlPFC connectivity was not significantly associated with the cognitive function index (*r* = −0.066, *p* = 0.44, 95% CI: [-0.23, 0.10]). Within-group analyses similarly showed no significant associations in older adults (*r* = 0.24, *p* = 0.059, 95% CI: [-0.01, 0.46]) or younger adults (*r* = −0.19, *p* = 0.114, 95% CI: [-0.40, 0.04]).

## Discussion

The present study investigated age-related differences in the LC activity and LC-PFC connectivity during absolute-ambiguity processing and their relationship with task dependent and task independent behavioral measures. Findings revealed heightened LC activity in older compared to younger participants in absolute-ambiguity condition. Notably, this higher LC activity in older adults was linked with slower RTs during the encoding of ambiguous expressions, indicating greater cognitive effort. This higher LC activation in older adults during ambiguity processing reflects increased cognitive effort, contributing to task-dependent behavioral response. Furthermore, older adults exhibited increased LC-dlPFC connectivity compared to younger adults, that associated with better task independent emotional resilience and mental wellbeing, indicating the importance of this pathway in supporting healthy aging.

### Positivity bias and better emotional resilience among older adults

Behaviorally, both age groups showed increased RTs when processing ambiguous faces, but only older adults exhibited a positivity bias, interpreting ambiguous faces as happy more frequently than fearful. This aligns with the socioemotional selectivity theory in aging (Carstensen & DeLiema, 2017). The positivity bias is consistent with findings in attention (Kim & Barber, 2022), memory (Charles et al., 2003; Ziaei et al., 2016), emotion recognition (Ziaei et al., 2015, 2016), and empathy (Ziaei et al., 2021). Older adults also demonstrated higher emotional resilience, suggesting their better mental well-being may reflect their tendency to interpret ambiguous signals more positively. Longer RTs for both groups indicate greater cognitive demands in processing ambiguous expressions (Kaminska et al., 2020), while lower confidence ratings during ambiguous trials highlight the task’s difficulty compared to unambiguous or intermediate conditions (Wang et al., 2017).

### Age-related enhanced LC activity during absolute-ambiguous condition

Neuroimaging results revealed elevated LC activity in older compared to younger adults during emotional ambiguity processing. This heightened LC activity in older adults associated with longer RTs, indicating greater cognitive effort in processing emotionally ambiguous stimuli. Prior studies have also shown that longer RTs reflect greater cognitive demands in conflict resolution tasks (Neto et al., 2021; Parr et al., 2023). Neurophysiological evidence indicates that LC activation scales with task difficulty, increasing with decision-related effort and longer RTs in complex tasks (Bornert & Bouret, 2021), reflecting sustained attention and cognitive effort (Rajkowski et al., 2004) underscoring the LC-NE system’s role in processing uncertainty and decision-making under ambiguity (Payzan-LeNestour et al., 2013). While Aston-Jones & Cohen (2005) linked high LC activity to faster RTs in simple, stimulus-driven tasks, this relationship shifts during effortful stimuli processing such as ambiguous faces. This indicates higher LC activation in older adults during ambiguity processing may reflect higher cognitive effort required for processing ambiguous signals at the cost of slower responses.

In the current study, the LC-NE system supports adaptive processes in healthy aging by enhancing task-relevant neural activity (Aston-Jones & Cohen, 2005). For example, the LC’s involvement in the emotional Stroop-task highlights its role in cognitive control and attentional flexibility (Grueschow et al., 2020). Recent studies have also highlighted higher LC activity among older adults during emotionally salient tasks, including reversal reinforcement learning (Ludwig et al., 2024) and during novel face encoding (Prokopiou et al., 2022). We extend these findings by showing that LC activity is selectively modulated by the absolute-ambiguity condition. This higher LC activity, compared to younger adults, enables older individuals to allocate additional cognitive resources to manage the heightened demands of processing ambiguous stimuli, highlighting the LC-NE system in supporting cognitive function during healthy aging. Additionally, LC activity was not elevated during other emotionally arousing conditions (fearful and happy expressions or the intermediate ambiguity conditions), suggesting specificity of the LC activity to process cognitively demanding stimuli, such as the absolute ambiguous condition in this study.

The observed elevated LC activity among older adults aligns with the Compensation-Related Utilization of Neural Circuits Hypothesis (CRUNCH), According to this hypothesis, older adults recruit additional neural resources to maintain cognitive functioning under increasing cognitive demands, compared to younger adults (Reuter-Lorenz & Cappell, 2008). Given that aging is accompanied by widespread changes in the brain’s structural and functional connections (Grady, 2012; Schulz et al., 2022; Kim & Kim, 2023), older adults may employ adaptive neural mechanisms to maintain attention and cognitive effort to resolve conflicting stimuli, in tasks such as processing emotional ambiguity.

While we observed elevated task-related LC activation in healthy older adults compared to younger adults, animal studies suggest aging alters tonic firing at rest, leading to reduced phasic responses due to their inverse relationship (Aston-Jones & Cohen, 2005; Kelberman et al., 2024). It also disrupts two key regulators of NE in aged mice: norepinephrine transporter (NET) which mediates NE reuptake, and α2a-adrenergic receptors, which provide inhibitory feedback—leading to prolonged and amplified LC neuronal activation (Budygin et al., 2024). However, further research is required to better understand how tonic and phasic LC activity operates in aging models during cognitively demanding or salient tasks.

The enhanced engagement of the LC-NE system also supports cognitive reserve, helping older adults sustain cognitive performance despite age-related neural changes (Stern, 2009; Mather, 2016). The NE released during cognitively demanding tasks provides neuroprotective benefits that help to mitigate inflammation and neurodegeneration in late life (Heneka et al., 2010). In the context of ambiguity processing, attention, cognitive effort, and exposure to novelty associated with emotional ambiguity processing might enhance the LC activation, ultimately supporting cognitive functions critical for resolving ambiguity in older adults. This aligns with studies showing that enhanced LC activity is essential for maintaining cognitive functions in healthy aging (Prokopiou et al., 2022; Ludwig et al., 2024). Previously, higher activations among older adults have been reported as a compensation for declining cognitive capacity (Aron et al., 2022). However, contrasting findings of no age-related difference in LC activity have also been noted (Berger et al., 2023). Conceptually, heightened LC activity in late adulthood may stem from region’s high metabolic demand, which makes it particularly susceptible to reactive oxygen species and toxins generated as byproducts of its activity with increasing age (Kelly et al., 2021; Matchett et al., 2021). These stressors can affect LC function, prompting the remaining neurons to compensate by increasing their activity to sustain cognition (Szot et al., 2016). Additionally, individual differences in vascular characteristics, such as in LC blood supply, may explain variability in the LC activation and cognitive performance among older adults (Bekar et al., 2012; Giorgi et al., 2020). Future studies are needed to further explore the mechanisms of LC neuronal compensation in LC, especially in healthy aging.

### Age-related increase in LC-dlPFC connectivity during ambiguity processing

Our results also revealed higher functional connectivity between the left LC and right dlPFC among older compared to younger adults during processing of absolute-ambiguous condition. This finding supports the hypothesis that processing emotional ambiguity becomes increasingly cognitively demanding with age (Liebherr et al., 2017; Gaubert et al., 2023) requiring additional neural connections to compensate for age-related cognitive decline. The role of dlPFC is well-established as a key region for cognitive effort and conflict resolution (Kane & Engle, 2002; Rahnev, 2017). Previous studies in aged monkeys have shown that the LC-NE system stabilizes dlPFC function through the phasic release of norepinephrine, enhancing LC-dlPFC connectivity and supporting working memory in aging (Arnsten et al., 2012; Cools & Arnsten, 2021). Consistent with this, our results suggest that stronger functional connectivity of the left LC with right dlPFC may support older adults to manage the heightened cognitive and emotional demands of processing ambiguous emotional expressions. This laterality likely reflects dlPFC functional specialization, with the right dlPFC being more attuned to visual affective content and the left dlPFC to processing verbal content (White et al., 2023). A potential limitation of gPPI analyses is the risk of inflated false positives with a liberal cluster-defining threshold (Eklund et al., 2016) which indicates the need for cautious interpretation.

### Higher LC-dlPFC connectivity supports mental wellbeing in aging

Importantly, left LC and right dlPFC connectivity was associated with better emotional wellbeing in older adults, evidenced by lower anxiety and stress scores, and higher emotional resilience. This suggests the LC-dlPFC pathway plays a crucial role in maintaining emotional stability with age. These results align with the autonomic compensation model that posits age-related NE hyperactivity and its connectivity with PFC aids emotional well-being in late life (Mather, 2024). Additionally, the dlPFC established role in stress and anxiety regulation (White et al., 2023), executive functions, such as cognitive control and emotional regulation (Chen et al., 2023; Nejati et al., 2021) supports this interpretation that the LC-dlPFC connectivity reflects top-down control mechanism, buffering against negative emotional states and promoting emotional resilience in later life. Therefore, our findings may suggest two complementary roles of the LC in healthy aging. During ambiguity processing, higher LC activation is linked to slower RTs, reflecting greater cognitive effort among older adults to perform the task. At the same time, stronger LC-dlPFC connectivity in late-life may serve as an adaptive function, regulating stress, anxiety and promoting emotional resilience. This suggests that while LC activation reflects heightened cognitive engagement, its connectivity with the dlPFC may enable top-down regulation, helping older adults manage cognitive and emotional demands.

Future research should examine whether the LC-dlPFC connection persists during stress-related tasks and its role in regulating emotional distress in real-life scenarios. Additionally, investigating the connections directionality could provide novel insights into the neural mechanisms underlying emotional regulation in aging. Lastly, no age-related differences were observed in the dmPFC, vmPFC, insula, ACC or FFA, underscoring the LC-dlPFC connection’s unique role in processing emotional ambiguity and supporting mental well-being in healthy aging.

## Limitations

This study has limitations. First, using static facial expressions may not fully reflect the complexity of real-life emotional processing or the ambiguity in emotional situations (Trautmann et al., 2009). Dynamic facial expressions could offer a more ecologically valid assessment of emotional ambiguity and better insights into LC activity during real-life processing. Second, the LC’s proximity to the 4th ventricle makes it susceptible to physiological artifacts, such as cardiac pulsatility and respiratory fluctuations. While aCompCor was used to regress noise from white matter and cerebrospinal fluid, future studies should include physiological measures to further minimize noise. Notably, our post-hoc analyses revealed no significant correlation between LC activity and the 4th ventricle, supporting the reliability of our findings.

In summary, our study highlights the pivotal role of LC activity and its connectivity with the dlPFC during emotional absolute-ambiguous task, and its link with better mental well-being in older adults. Our findings revealed heightened LC activity and stronger LC-dlPFC connectivity in older adults, compared to younger individuals, highlighting the importance of this neural pathway in managing cognitively demanding tasks while maintaining emotional wellbeing in healthy aging. By emphasizing the significance of this functional connectivity in promoting healthy aging, our results open avenues for future research into targeted interventions aimed at enhancing cognitive functions, and emotional well-being in late adulthood.

## Footnotes

- We would like to thank radiographers and MR physicists at the 7T MR center at NTNU for their help during this project. We also would like to thank our participants for their time and effort during the experiment. We thank Jørgen Østmo-Sæter Olsnes, Stian Framvik, Avneesh Jain, Jae Hong, and Karina Tømmerdal for their help during data collection. This project was funded by Kavli Foundation and was partially supported by the Research Council of Norway through its Centers of Excellence scheme, project number 332640.
- The authors declare no competing financial interests
- Correspondence should be addressed to Maryam Ziaei, and Arjun Dave
- Data availability: Behavioral data, ROI signals and the stimuli can be found in the Open Science Framework from this link: https://osf.io/7v2ez

---

**Figure S1: Emotional Ambiguity Processing Across Age groups.** Statistical parametric map depicting whole-group activation during the absolute-ambiguity processing condition. Additionally, the right and left dlPFC are marked with circles, highlighting their involvement in ambiguity processing. Activation maps are displayed after multiple corrections at *P_FDR_* < 0.05, overlaid on a standard MNI template.

**Figure S2:** Overlap between observed BOLD activation and the LC mask with small volume correction (SVC) confined within the LC mask. The white line outlines the LC mask, while red highlights activated voxels (inside vs outside mask) after the SVC. The included cluster represents significant voxels retained through SVC.

**Figure S3: Brainstem Activation Across Participants During Absolute Ambiguity Condition.** Whole-brainstem activation map thresholded at p < 0.005 uncorrected with a voxel extent threshold of ≥10 voxels during the absolute ambiguity condition across the participants.

**Figure S4: Emotional Ambiguity Processing Between Age Groups.** Statistical parametric maps showing group differences in activation during the absolute-ambiguity condition (Younger adults > Older adults, and Older adults > Younger adults). Activation maps are displayed after multiple corrections at *P_FDR_* < 0.05, overlaid on a standard MNI template.

