## Supplemental Tables for "Age-related Increase in Locus Coeruleus Activity and Connectivity with Prefrontal Cortex during Ambiguity Processing"

**Supplementary/Extended Tables**

**Table S1:** Participant Demographics, Cognitive Behavioral Task Performance, and Self-Reported Measures for Younger and Older Adults

| **Measurement** | **Younger adults**  ***Mean SD*** | | **Older adults**  ***Mean SD*** | | **Older>Younger**  ***Independent t-test*** |
| --- | --- | --- | --- | --- | --- |
| ***Demographic*** |  |  |  |  |  |
| Age | 25.80 | 4.02 | 71.30 | 4.11 | 67.59** |
| Gender | 40M, 35F | - | 34M,35F | - | - |
| Years of Education | 16.79 | 4.11 | 12.05 | 4.62 | -6.43** |
| ***Cognition*** |  |  |  |  |  |
| Trail Making (B-A) | 26.67 | 15.62 | 42.63 | 41.75 | 3.01* |
| Raven’s Progressive Matrices | 0.77 | 0.16 | 0.49 | 0.26 | -8.60** |
| Digit Span - forward | 6.15 | 1.32 | 5.23 | 1.38 | -4.06** |
| Digit Span - backward | 5.19 | 1.14 | 3.91 | 1.17 | -6.62** |
| Stroop effect | 0.21 | 0.15 | 0.31 | 0.16 | 3.90** |
| COWAT - Phonemic | 12.60 | 3.68 | 15.50 | 4.56 | 4.56* |
| COWAT - Semantic | 15.57 | 4.06 | 15.66 | 3.58 | -0.40 |
| RMET | 0.66 | 0.12 | 0.60 | 0.11 | -3.26** |
| ***Well-being measurements*** |  |  |  |  |  |
| DASS - Depression | 9.97 | 9.17 | 2.43 | 3.12 | -6.49** |
| DASS – Anxiety | 8.72 | 6.48 | 2.14 | 2.84 | -7.77** |
| DASS - Stress | 13.23 | 7.99 | 5.33 | 5.66 | -7.36** |
| DERS | 80.59 | 17.44 | 67.03 | 9.58 | -5.71** |
| GHQ28 – somatic symptoms | 13.00 | 3.46 | 10.32 | 2.07 | -5.58** |
| GHQ28 - Anxiety/Insomnia | 12.93 | 3.86 | 10.33 | 2.52 | -4.74** |
| GHQ28 – Social Dysfunction | 13.79 | 2.89 | 13.26 | 1.57 | -1.34 |
| GHQ28 – Severe Depression | 10.25 | 3.99 | 7.51 | 1.22 | -5.48** |
| HADS – Depression | 4.51 | 3.58 | 1.46 | 1.47 | -6.57** |
| HADS – Anxiety | 6.75 | 3.51 | 2.88 | 2.33 | -7.70** |
| CD-RISC | 68.97 | 14.33 | 76.62 | 9.28 | 3.77** |
| PSS | 1.63 | 0.65 | 1.05 | 0.38 | -6.30** |
| IUS | 59.08 | 15.90 | 46.61 | 14.18 | -4.95** |
| IRI Cognitive empathy | 34.56 | 8.84 | 32.01 | 6.92 | -1.91 |
| IRI Perspective Taking | 18.61 | 4.33 | 18.78 | 3.78 | 0.22 |
| IRI Fantasy Scale | 15.94 | 6.44 | 13.24 | 5.13 | -2.79** |
| IRI Emotional empathy | 30.50 | 7.73 | 31.94 | 6.62 | 1.21 |
| IRI Empathic concern scale | 18.98 | 4.77 | 20.42 | 4.32 | 1.91 |
| IRI Personal distress scale | 11.53 | 5.19 | 11.52 | 4.15 | -0.01 |
| STAI-state anxiety | 34.92 | 10.20 | 27.8 | 6.30 | -5.19** |
| STAI-trait anxiety | 35.53 | 9.30 | 25.0 | 5.20 | -8.71** |
| ***PCA Composite Scores*** |  |  |  |  |  |
| Emotional Resilience Index | -0.54 | 1.03 | 0.60 | 0.49 | -8.32** |
| Cognitive Function Index | 0.45 | 0.71 | -0.49 | 1.03 | 6.38** |
| ***Task performance***  ***RT (milliseconds)*** |  |  |  |  |  |
| Anchor | 0.60 | 0.10 | 0.63 | 0.11 | -0.24 |
| Intermediate Ambiguity | 0.69 | 0.12 | 0.68 | 0.14 | 0.09 |
| Absolute-Ambiguity | 0.75 | 0.14 | 0.72 | 0.20 | 0.12 |
| ***Response bias*** |  |  |  |  |  |
| Ratio of happy responses | 49.45 | 6.03 | 53.08 | 6.05 | -3.60** |
| ***Confidence rating*** |  |  |  |  |  |
| Anchor | 1.60 | 0.69 | 1.76 | 0.73 | -1.32 |
| Intermediate | 1.94 | 0.81 | 2.23 | 0.71 | -2.33* |
| Absolute-Ambiguity | 1.37 | 0.65 | 1.51 | 0.70 | -1.22 |

*Note: Stroop Effect (a measure of cognitive interference measured by response time in incongruent trials – response time in neutral trials)/response time in neutral trials); COWAT (Controlled Oral Word Association Test); RMET (Reading the Mind in the Eyes Test); DASS (Depression Anxiety Stress Scales); DERS (Difficulties in Emotion Regulation Scale; GHQ (General Health Questionnaire), HADS (Hospital Anxiety and Depression Scale); CD-RISC (Connor-Davidson Resilience Scale), PSS (Perceived Stress Scale); IUS (Intolerance of Uncertainty Scale); IRI (Interpersonal Reactivity Index); STAI (State-Trait Anxiety Inventory); PCA (Principal component Analysis);*p<0.05, **p<0.001.*

**Table S2:** Peak coordinates of activated clusters for absolute ambiguity condition across participants in brainstem

| **Hemisphere** | **MNI coordinates**  **X Y Z** | **Cluster Size** | **Cluster level**  **P_UNC_** | **Cluster level**  **P_FDR_** | **Peak level**  **P_UNC_** | **Peak level**  **P_FDR_** | ***t*-value** | ***Labels*** |
| --- | --- | --- | --- | --- | --- | --- | --- | --- |
| R | [12 -40 -44] | 77 | <0.001 | 0.054 | <0.001 | <0.001 | 5.83 | Caudal Medulla |
| L | [-8 -28 -30] | 20 | 0.045 | 0.498 | <0.001 | 0.008 | 4.60 | Mid-to-Caudal Pons |
| L | [-8 -20 -34] | 101 | <0.001 | 0.007 | <0.001 | 0.034 | 4.14 | Rostral Pons |
| L | [14 -28 -36] | 14 | 0.087 | 0.657 | <0.001 | 0.036 | 4.13 | Caudal Pons |
| L | [-8 -38 -50] | 21 | 0.041 | 0.498 | <0.001 | 0.050 | 4.01 | Caudal Medulla |

***MNI*** *= Montreal Neurological Institute;* ***L*** *= left;* ***R*** *= right;* **P_UNC_** *= P_UNCORRECTED_;* **P_FDR_** *= P_FALSE DISCOVERY RATE_; cluster size ≥ 10 voxels*
