## Supplemental Text for "Age-related Increase in Locus Coeruleus Activity and Connectivity with Prefrontal Cortex during Ambiguity Processing"

**Supplementary Text/Results**

**Locus Coeruleus (LC) Volume**

LC ROIs for each participant were derived by warping the template-based LC mask to subject space, yielding volumes within the expected range reported in prior studies (Langley et al., 2016; Castellanos et al., 2015; Schwarz et al., 2016). The Left LC volume was 16.06 ± 2.31 mm³ in the young group and 15.81 ± 2.26 mm³ in the old group, with no significant difference (*t* = 0.43, p = 0.66). Similarly, the Right LC volume was 15.07 ± 2.64 mm³ in the young group and 15.30 ± 2.67 mm³ in the old group, also showing no significant difference (*t* = -0.35, p = 0.72). These findings indicate that LC volume does not significantly differ between age groups, further supporting the validity of the segmentation approach.

While we report LC volume estimates for descriptive purposes and for assessing its alignment with expected values from prior neuroimaging studies—we would like to point out that these methods consistently underestimate true LC volume. This underestimation is due to limited spatial resolution, partial volume effects (Engels-Domínguez et al., 2023), and the challenge of capturing the full caudal extent of the LC (Van Egroo et al., 2021). Therefore, these estimates should not be interpreted as biologically precise markers of LC size.
