## Supplementary figures and images for "Age-related Increase in Locus Coeruleus Activity and Connectivity with Prefrontal Cortex during Ambiguity Processing"

### Supplemental Figure 1

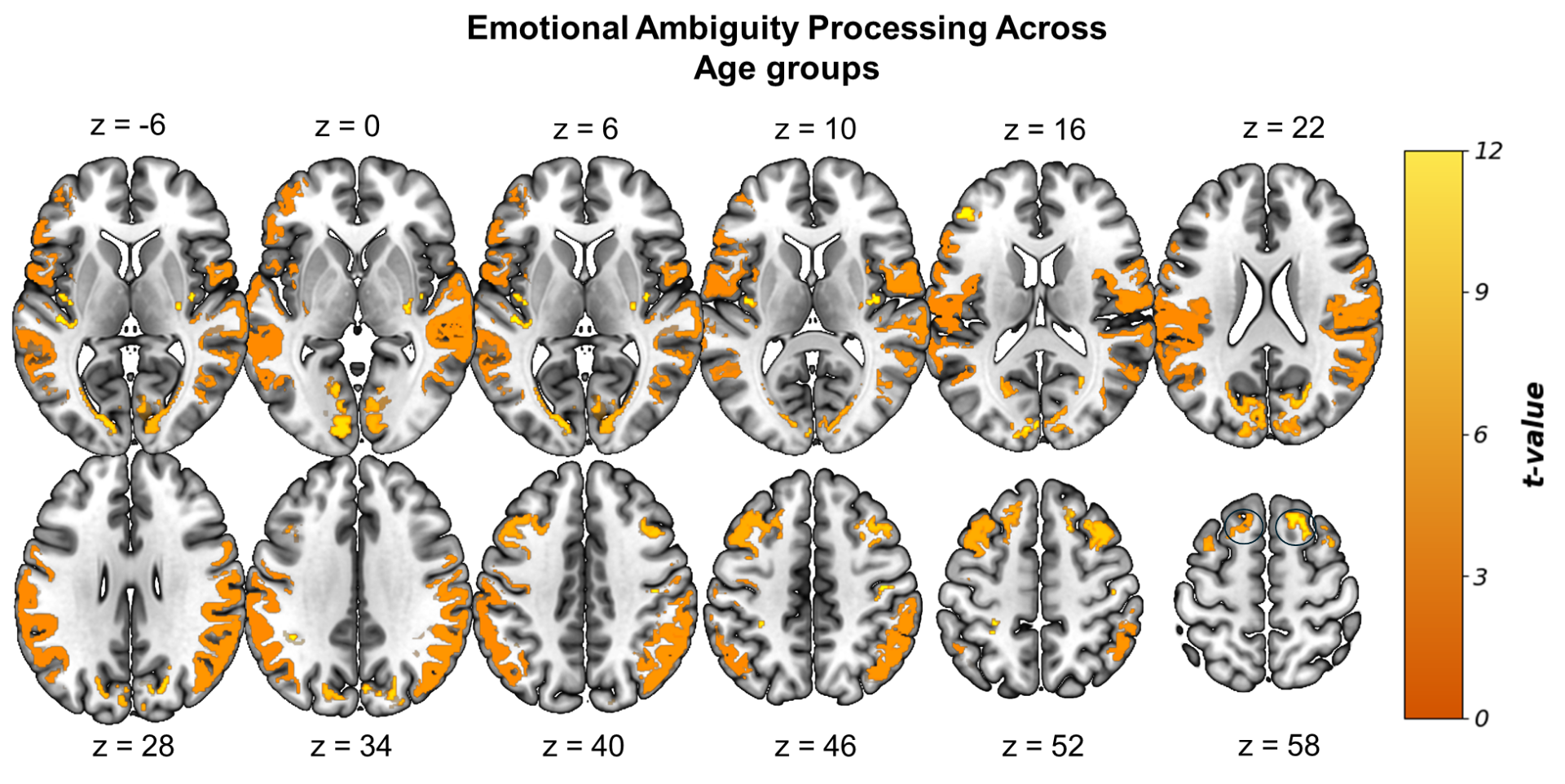

### Supplemental Figure 2

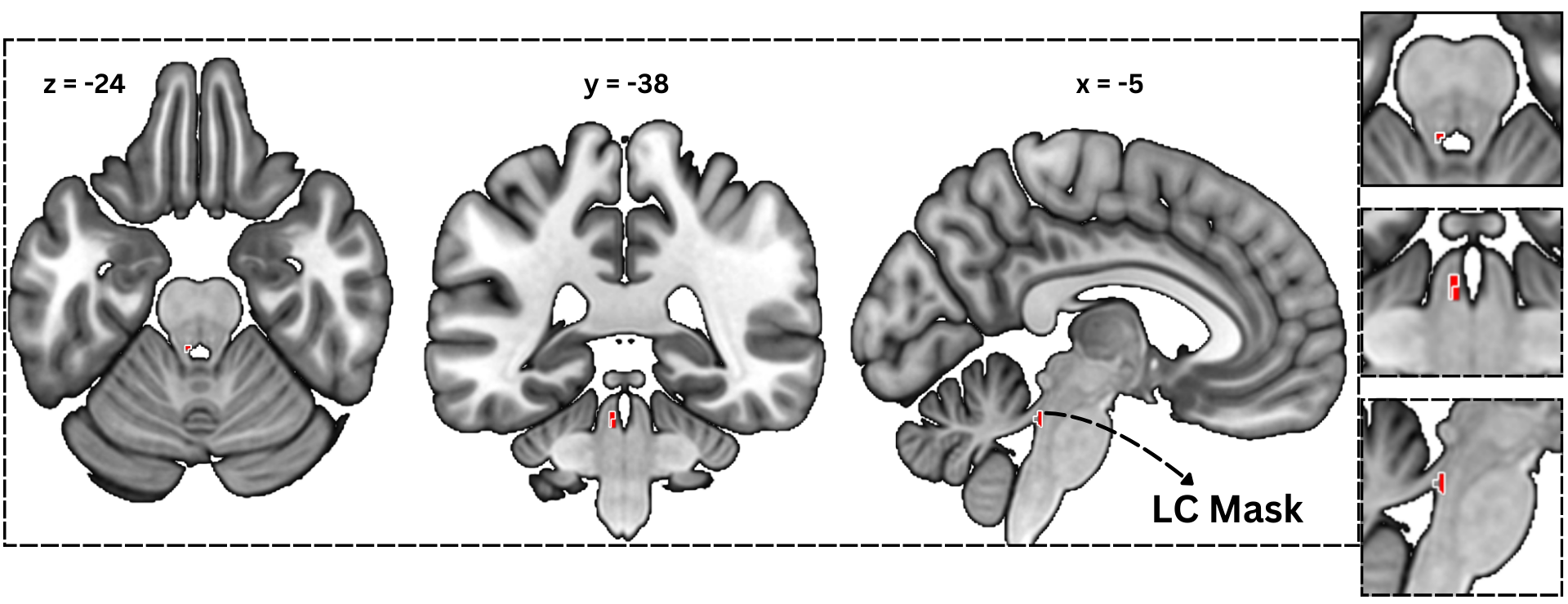

### Supplemental Figure 3

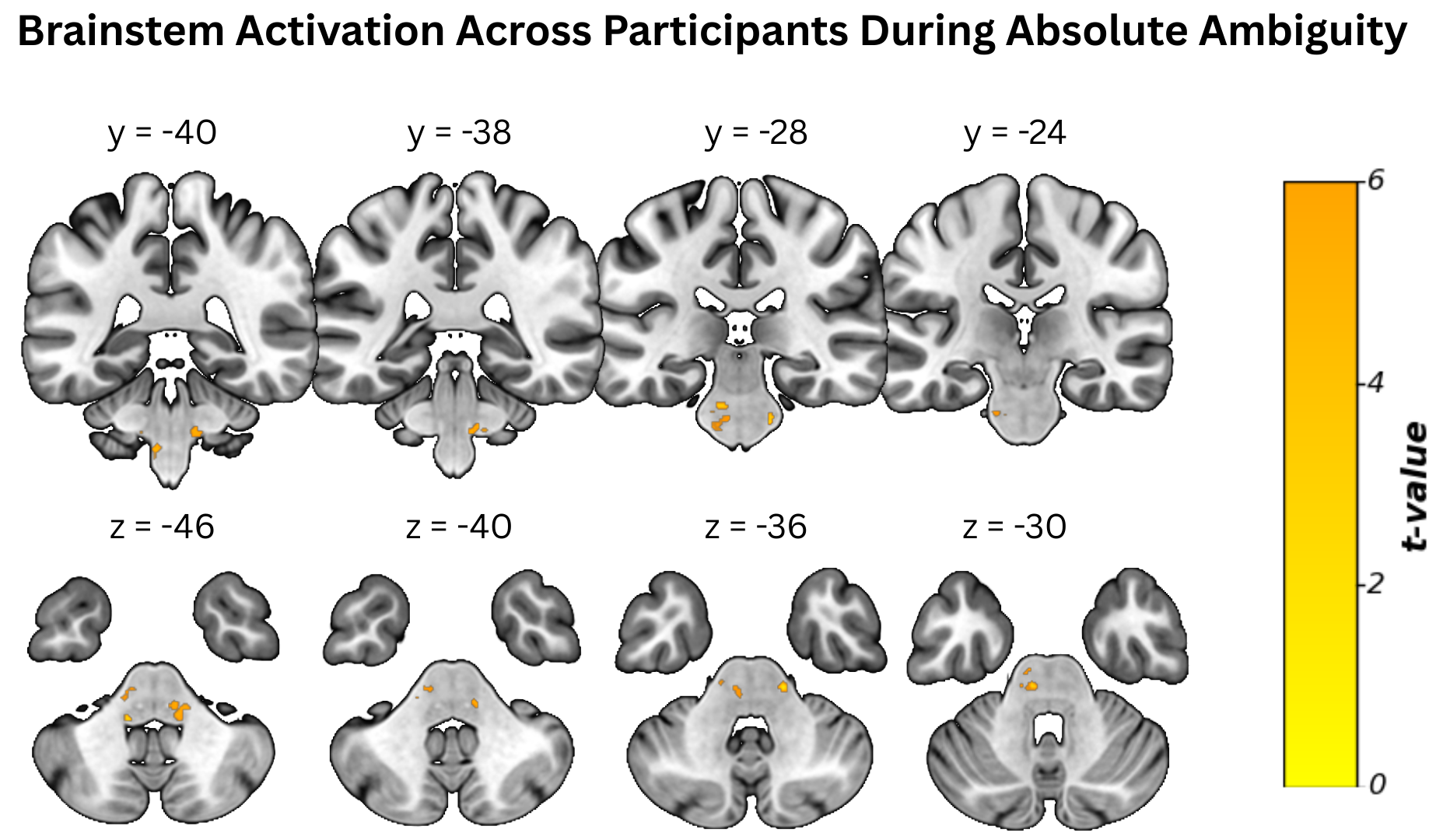

### Supplemental Figure 4

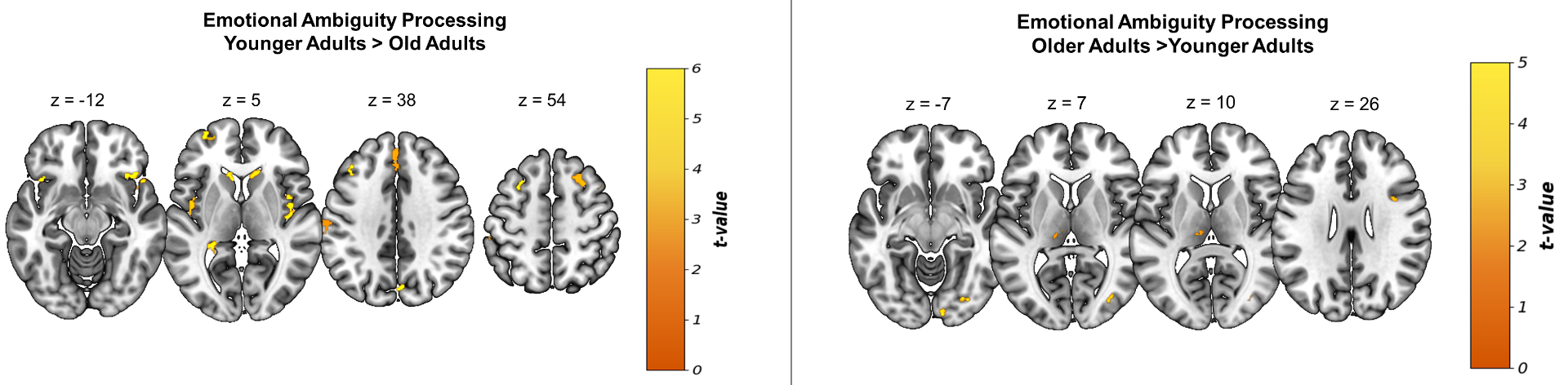
